# Simultaneous mesoscopic Ca^2+^ imaging and fMRI: Neuroimaging spanning spatiotemporal scales

**DOI:** 10.1101/464305

**Authors:** Evelyn MR Lake, Xinxin Ge, Xilin Shen, Peter Herman, Fahmeed Hyder, Jessica A Cardin, Michael J Higley, Dustin Scheinost, Xenophon Papademetris, Michael C Crair, R Todd Constable

**Author notes:** Equally contributing co-first author. Jointly supervised the research (Michael C Crair, R Todd Constable).

## Abstract

To achieve a more comprehensive understanding of brain function requires simultaneous measurement of activity across a range of spatiotemporal scales. However, the appropriate tools to perform such studies are largely unavailable. Here, we present a novel approach for concurrent wide-field optical and functional magnetic resonance imaging (fMRI). By merging these two modalities, we are for the first time able to simultaneously acquire whole-brain blood-oxygen-level-dependent and whole-cortex calcium-sensitive fluorescent measures of brain activity. We describe the developments that allow us to combine these modalities without compromising the fidelity of either technique. In a transgenic murine model, we examine correspondences between activity measured using these modalities and identify unique and complementary features of each. Our approach links cell-type specific optical measurements of neural activity to the most widely used method for assessing human brain function. These data and approach directly establish the neural basis for the macroscopic connectivity patterns observed with fMRI.

## INTRODUCTION

At a range of spatiotemporal scales, from individual cells to local circuits to large-scale networks, a wide array of tools has been developed to examine the organizational principles of the nervous system.^1–4^ For example, the advent of genetically encoded calcium (Ca^2+^) indicators enabled researchers to image cell-specific activity and investigate the correlations between this activity, local circuit dynamics, and behavior.^5–8^ Recently, approaches for carrying out wide-field “mesoscopic” imaging of the entire mouse neocortical surface have significantly broadened our understanding of coordinated signaling between distant areas.^9–12^ These signals are thought to largely reflect Ca^2+^ influx resulting from somatic action potentials, thereby providing high spatiotemporal resolution measurements of neural activity.^13–14^ These indicators can be targeted to genetically specified cell types, but necessarily require invasive manipulation of the nervous system and are limited to non-human models.

In contrast, functional magnetic resonance imaging (fMRI) provides a completely non-invasive means of measuring activity in both human and non-human subjects throughout the whole brain, including deep structures. However, fMRI has limited spatiotemporal resolution and relies on a complex indirect measure of neural activity through a blood oxygen level-dependent (BOLD) contrast mechanism. Few studies have compared fMRI with simultaneous data from other modalities that directly measure neural activity, and most of these studies are limited to a single region of interest (ROI) and task-based evoked responses.^15–18^ Given the whole-brain nature of this intrinsic contrast mechanism associated with brain activity, and the ability to perform studies in both animals and humans, research in fMRI has rapidly expanded to the point where there are tens of thousands of peer reviewed publications per year that use this modality.

Despite the considerable success of these approaches individually, technical challenges to simultaneous monitoring of brain activity across spatiotemporal scales limits our ability to understand how large-scale functional networks emerge from local populations of synaptically connected neurons. Meeting this challenge requires tools that enable the simultaneous collection of multi-modal imaging data. Here, we present a novel system capable of performing simultaneous mesoscopic Ca^2+^ imaging of neural activity and fMRI in mice.

Our results provide the first independent validation of the neural source of functional connectivity measured with fMRI in the resting-state. We demonstrate that sensory inputs drive similar average signals across modalities when considering the different spatiotemporal response functions in each. We also show that trial-to-trial variations in response magnitude are correlated across modalities, indicating a common neural source for these fluctuations. Finally, we demonstrate that functional parcellation of the cortex based on connectivity derived from measurements of spontaneous activity are highly consistent across modalities. Together, these results provide critical validation of functional connectivity as measured with fMRI and demonstrate an important new approach to compare cellular activity in genetically targeted cells with BOLD signals, linking cell specific activity in rodents with the macroscopic circuits examined in animal models and humans.

## RESULTS

Outside of the MRI environment, wide-field Ca^2+^ imaging is typically performed with a conventional microscope and camera positioned directly above the animal. To enable multi-modal dual-imaging, we redesigned the imaging apparatus to work within the confined space and high magnetic field of the MRI scanner. This included the development of both customized optics and radio frequency (RF) MRI hardware. In the following sections, we describe the novel hardware and software developments necessary for simultaneous multi-modal data acquisition and analysis.

### Simultaneous Ca^2+^ and MR-imaging

A schematic of the experimental set-up is shown in Fig. 1.a. Calcium imaging was performed through the intact skull on transgenic mice expressing GCaMP6f in cortical excitatory neurons (Slc17a7-Cre/Camk2a-tTA/Ai93 mice, **Methods**).^19–22^ However, it should be emphasized that the methods described here are suitable for any mouse line with a sufficiently bright optical signal (we have tested: Emx/Camk/TeTG6f; Slc17a7/Camk/TeTG6f; Slc17a7/Ai162; Vip/Ai162, and Som/Ai162 lines, data not shown).

**Fig. 1.**
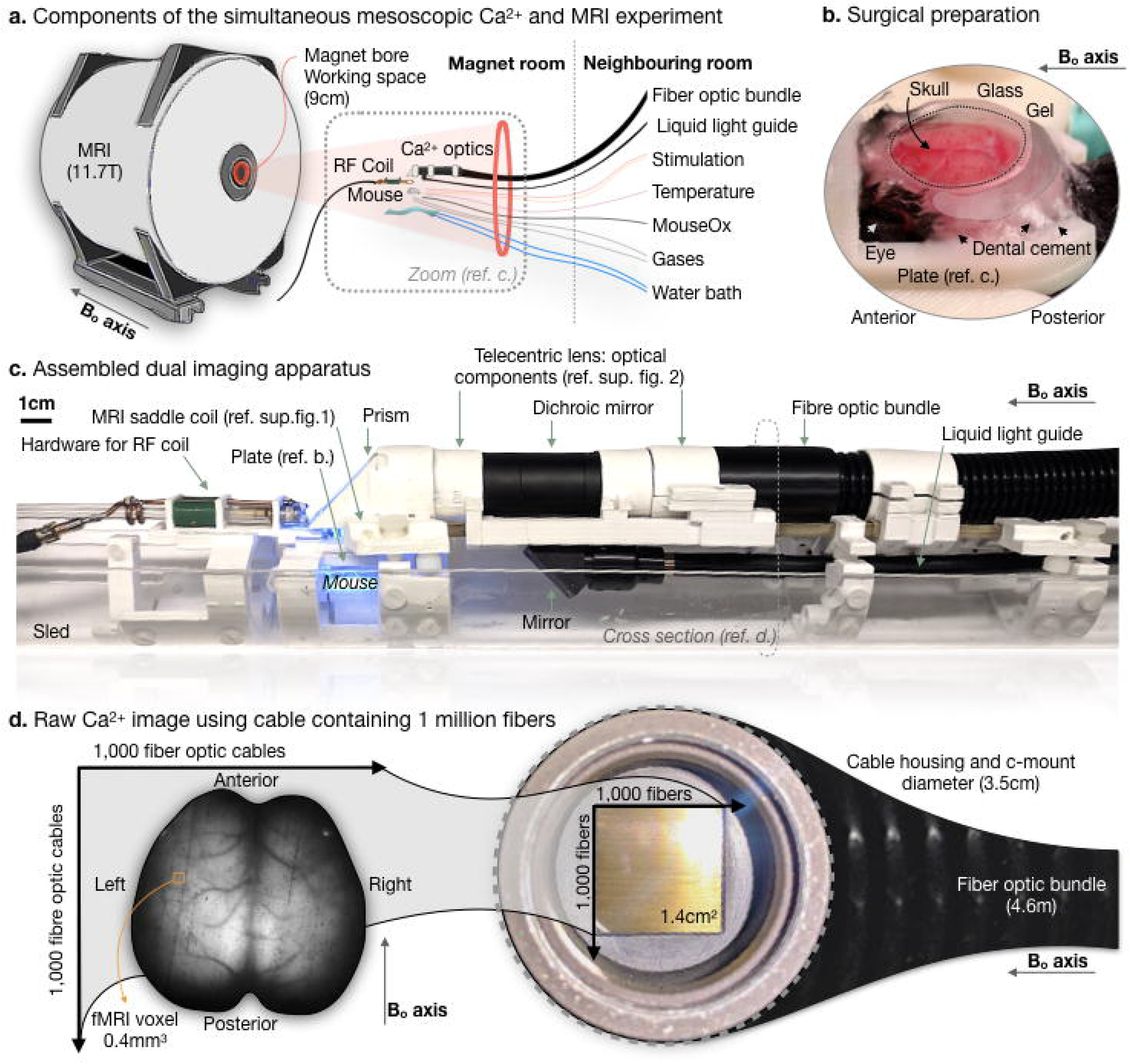
Simultaneous mesoscopic Ca^2+^ and MR-imaging experimental setup. An overview of the components (a.), which include an 11.7T. 9cm bore MR-scanner, into which we insert the optical components for Ca^2+^ signal acquisition, MRI coil, animal, and physiological monitoring and maintenance equipment. The Ca^2+^ signal is collected through an intact skull. Thus, a minimally invasive surgical preparation is necessary, and a prepared animal is shown in (b.). Following resection of the scalp and tissue overlying the skull surface, a ‘well’ of dental cement is created around the circumference of the cleaned skull. The well is filled with optically transparent Phytagel and sealed with a glass cover-slip. This minimizes susceptibility artifacts in MR-images and provides a smooth surface that facilitates Ca^2+^ signal transmission. The preparation is secured to a plate that cradles the sides of the skull and attaches to the skull above the olfactory bulb. The plate is fixed to the MRI coil below the Ca^2+^ imaging hardware. A photograph of the assembled dual-imaging apparatus is shown in (c.). The Ca^2+^ imaging data is recorded by a CCD camera connected to a computer housed in a room adjacent to the. A 4.6m long fiber optic bundle relays the Ca^2+^ signal from within the magnet to the CCD camera. A raw Ca^2+^ image captured using this set-up is shown in (d.). A fiber optic bundle containing ~2 million fibers is used to obtain a FOV spanning the optically exposed FOV.

The surgical preparation was optimized for dual-imaging (**Methods**) to reduce fMRI signal loss caused by nearby materials with different magnetic susceptibilities, such as brain and air, without compromising fluorescent signal transmission. In a brief and minimally invasive surgical preparation, a dental cement ‘well’ was built along the circumference of the cleaned skull surface, filled with an MRI compatible optically transparent gel, and sealed with a glass cover-slip (Fig. 1.b). The gel allows the skull to be optically exposed for Ca^2+^ imaging, while minimizing the susceptibility effect in the fMRI acquisition. The skull was fixed to a custom holder that dovetails with an in-house built RF MRI coil and the optical imaging apparatus (Fig. 1.c). The RF coil was shaped like a saddle to avoid obstructing the optical field of view (FOV) and to optimize whole-brain fMRI-sensitivity (**Supplementary Fig. 1**).

Optical imaging was performed with a heavily modified telecentric lens. The primary modification was to remove all metal parts from the lens and replace them with nonconducting plastic (**Supplementary Fig. 2**). The coaxial illumination was also modified to accommodate fluorescent imaging. A key feature of the optical setup was a custom coherent fiber-optic bundle containing nearly two million elements that relayed the fluorescent signals from within the scanner to a camera housed in an adjacent room away from the high magnetic field (Fig. 1.d). The optical imaging device presented here is quite different from previous implementations combining MRI with simultaneous fluorescent Ca^2+^ data collection, where typically only single fibers were used to measure the fluorescence as a function of time from one or more ROIs.^23–28^ Here, we obtained dynamic high-resolution movies (FOV: 14.5×14.5mm^2^, resolution: 25×25μm^2^) of Ca^2+^ fluorescence activity spanning a large fraction of the mouse cortex. The combined dual-imaging set-up was assembled within a plastic sled that was inserted and secured within an 11.7T preclinical Bruker magnet (Bruker, Billerica, MA). This design can be easily scaled for larger animals, different bore sizes, and different magnetic field strengths.

### Multi-modal acquisition and image registration

We synchronously recorded optical and fMR images (Fig. 2.a) together with physiological data (**Supplementary Fig. 3**). Each fMRI volume (TR=1000ms, 1Hz, and spatial resolution 0.4×0.4×0.4mm^3^; see **Methods**) triggered the capture of 20 optical frames (20Hz). Fluorescence and fMRI data were processed using standard procedures, including motion correction, bandpass filtering, drift correction and global signal regression (**Supplementary Fig. 4** and **Supplementary Fig. 5**; and see **Methods**).

**Fig. 2.**
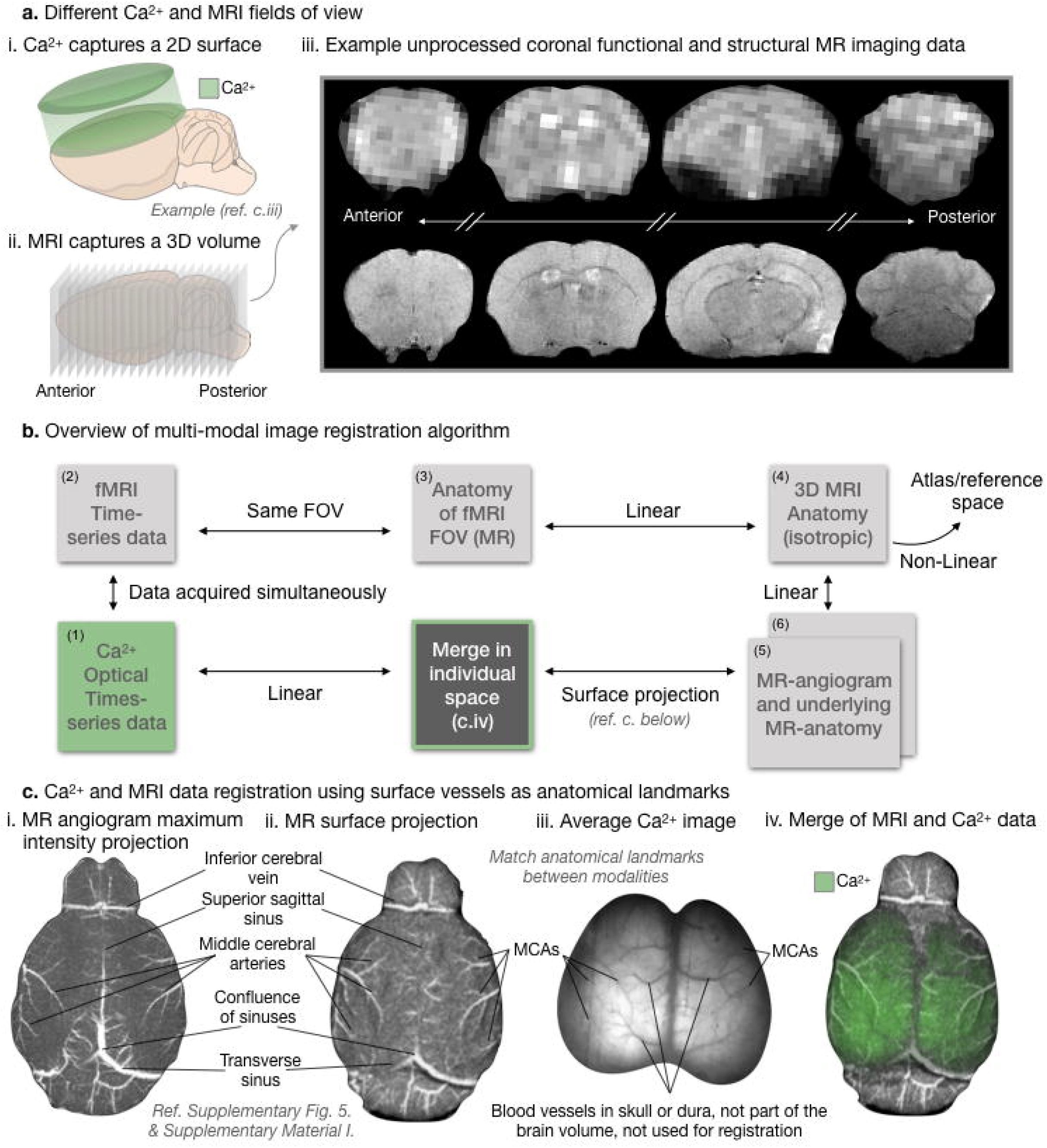
Registration pipeline for simultaneously acquired Ca^2+^ and MRI data. The registration of Ca^2+^ and fMRI data is non-trivial because the Ca^2+^ data is a 2D projection of the cortical surface (tissue depth contributing to the signal is unknown), and the fMRI data spans a 3D volume. Cartoons depicting the Ca^2+^ (i.) and functional MRI (ii.) acquisitions are shown in (a.). Sample MR functional (top row) and structural (bottom row) images are shown in (a.iii.). For multi-modal image registration (b.), in addition to the Ca^2+^ (1) and fMRI (2) simultaneously acquired data, we acquire: a high in-plane resolution 2D image of the fMRI FOV (3), an isotropic 3D anatomical MRI of the whole brain (4). an MR-angiogram (5). and a high-resolution anatomical image of the tissue within the angiogram FOV (6). First, using a rigid registration algorithm, the functional and structural MRI data are registered in ‘individual space’, and then non-linear registration is used to register each individual mouse to an atlas or reference space. To register the Ca^2+^ and MRI data (c.). we begin by using a custom ray-casting algorithm (**Supplementary Fig. 6. and Supplementary Material**) on the MR-angiogram data (c.i.) to create a surface projection (c.ii.). Surface vessels (e.g. projections from the middle cerebral arteries) and anatomical features (e.g. midline) visible in the MR-surface projection and the Ca^2+^ images (c.iii.) are used to move the Ca^2+^ and MRI-data to the same space (merged image, c.iv.).

Co-registration of the simultaneously recorded Ca^2+^ (2D-surface) and fMR (3D-volume) images was a challenging image registration task (Fig. 2.b). This was accomplished using the anatomy of the vasculature on the surface of the cortex, which is visible in both the optical images and in an MR-angiogram, as reference landmarks. Neither imaging modality requires the administration of an exogenous contrast agent.

In addition to the Ca^2+^ and fMRI time-series data, multi-modal image registration used an averaged Ca^2+^ image, a high in-plane resolution anatomical MR image that matched the fMRI scan prescription, an isotropic high-resolution MR anatomical image of the whole-brain, an MR-angiogram, and a high-resolution image of the brain tissue within the MR-angiogram FOV (see Fig. 2.b, Fig. 2.c, and see **Methods**). These latter two images were combined to make one MR surface projection image (2D) that recapitulates what the surface of the mouse brain would look like from above using MRI data (Fig. 2.c**, Supplementary Fig. 6**, and Methods).

Briefly, the high in-plane resolution anatomical images that match the fMRI scan prescription were first registered to the fMRI data and to the isotropic high-resolution anatomical image using a rigid registration algorithm that employs the normalized mutual information metric.^32^ Next, the angiogram and the associated anatomic image were registered to the isotropic high-resolution whole-brain image. These images in turn were used to generate the MR surface projection image. These steps move all the MRI data into the ‘animal’s reference space’. Using the isotropic high-resolution anatomical image, the data from each animal can be moved to an atlas (such as the The Allen Brain Atlas http://www.brain-map.org) or any other reference space using a non-linear, non-rigid registration method again based on normalized mutual information.^33,34^ Finally, the average Ca^2+^ image was registered to the MR surface projection image, which brings all the multi-modal data into the same space. All image registration was performed using BioImageSuite Web (www.bioimagesuite.org). Modules within which were custom developed in-house for this purpose and are freely available online.

### Evoked multi-modal Ca^2+^ and fMRI responses

We first demonstrate the utility of this multi-modal imaging approach with evoked responses to unilateral hind-paw electrical stimulation (Fig. 3). Using a data-driven approach, we identified the responding Ca^2+^ and fMRI ROIs (generalized linear modeling, **Supplementary Fig. 7**, and **Methods**), and observed robust evoked responses from both modalities in overlapping regions of cortex (**Supplementary Fig. 8**).

**Fig. 3.**
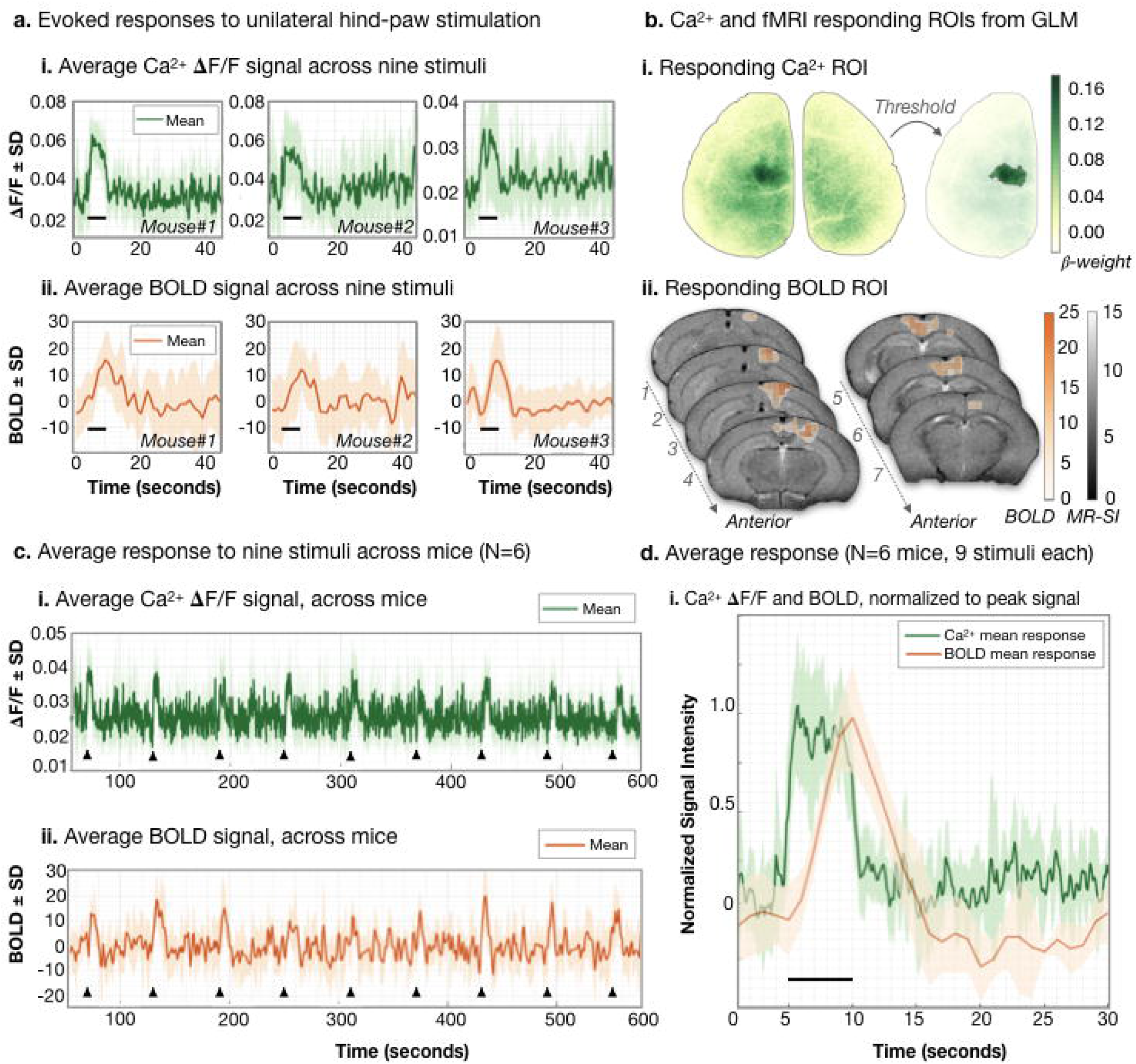
Responses to unilateral hind-paw stimulation. The ROIs for Ca^2+^ and fMRI responses to unilateral hind-paw stimulation were determined using standard task-based general linear modeling (Methods). The average responses (to N=9 stimuli) for three mice are plotted in (a.). The stimulus ON time is denoted by a black line in (a. and d.). The Ca^2+^ data (green) is plotted in the top row (i.). and corresponding fMRI data (orange) is plotted in the bottom row (ii.). The fMRI signal is normalized to the mean. In all plots, the standard deviation across stimulus presentations (or across mice) is shaded. Ca^2+^ and fMRI ROI data for Mouse#2 is shown in (b.). All unilateral hind-paw imaging experiments were 10 minutes (600 seconds) long and included 10 stimuli. The first response was excluded from all analyses to avoid the Ca^2+^ baseline drift (due to photobleaching) which significantly affected the first response in all recordings. Plotted in (c.) are the average responses across N=6 mice (N=9 stimuli from each mouse). Stimulus onset in (c.) is denoted by black triangles. The average, normalized (to peak amplitude) response across N=6 mice. N=9 stimuli each, is plotted in (d.) and this illustrates the different temporal dynamics of these two modalities. The fMRI signal is delayed, relative to stimulus presentation, as it is dependent in part on blood flow effects which are slow. Because of the complex relationship between blood flow, volume, and oxygenation, there is typically an undershoot observed after the stimulus is turned off in fMRI evoked responses.

To examine the relationship between the magnitude of the Ca^2+^ and fMRI evoked responses, hind-paw stimuli were applied at a constant current and frequency. Individual response amplitudes were quantified as the averaged signal within the responding ROI from the peak of the response until three seconds after the peak. The peak of the response was defined using the time-to-peak (TTP), calculated from one run (9 stimuli) for each of six mice (Ca^2+^ TTP: 0.6 ± 0.03 seconds, and fMRI TTP: 4.7 ± 0.5 seconds, mean ± standard deviation, SD, **Supplementary Fig. 9**). Previous work demonstrates that changing stimulus parameters (strength, duration) modulates the magnitude of Ca^2+^ and fMRI responses.^23^ We observed that the fluctuations in Ca^2+^ and fMRI evoked responses to individual identical stimuli were moderately but significantly correlated (Pearson’s correlation, r=0.28 P<0.04, Fig. 4). This extends previous work by demonstrating stimulus-by-stimulus spontaneous multi-modal co-variation in response amplitude. This attests to the high quality of both the Ca^2+^ and fMRI data collected, and because the measures report different sources of contrast, provides the first demonstration that the small spontaneous fluctuations in individual responses to identical stimuli are not noise, but are actually biologically meaningful.

**Fig. 4.**
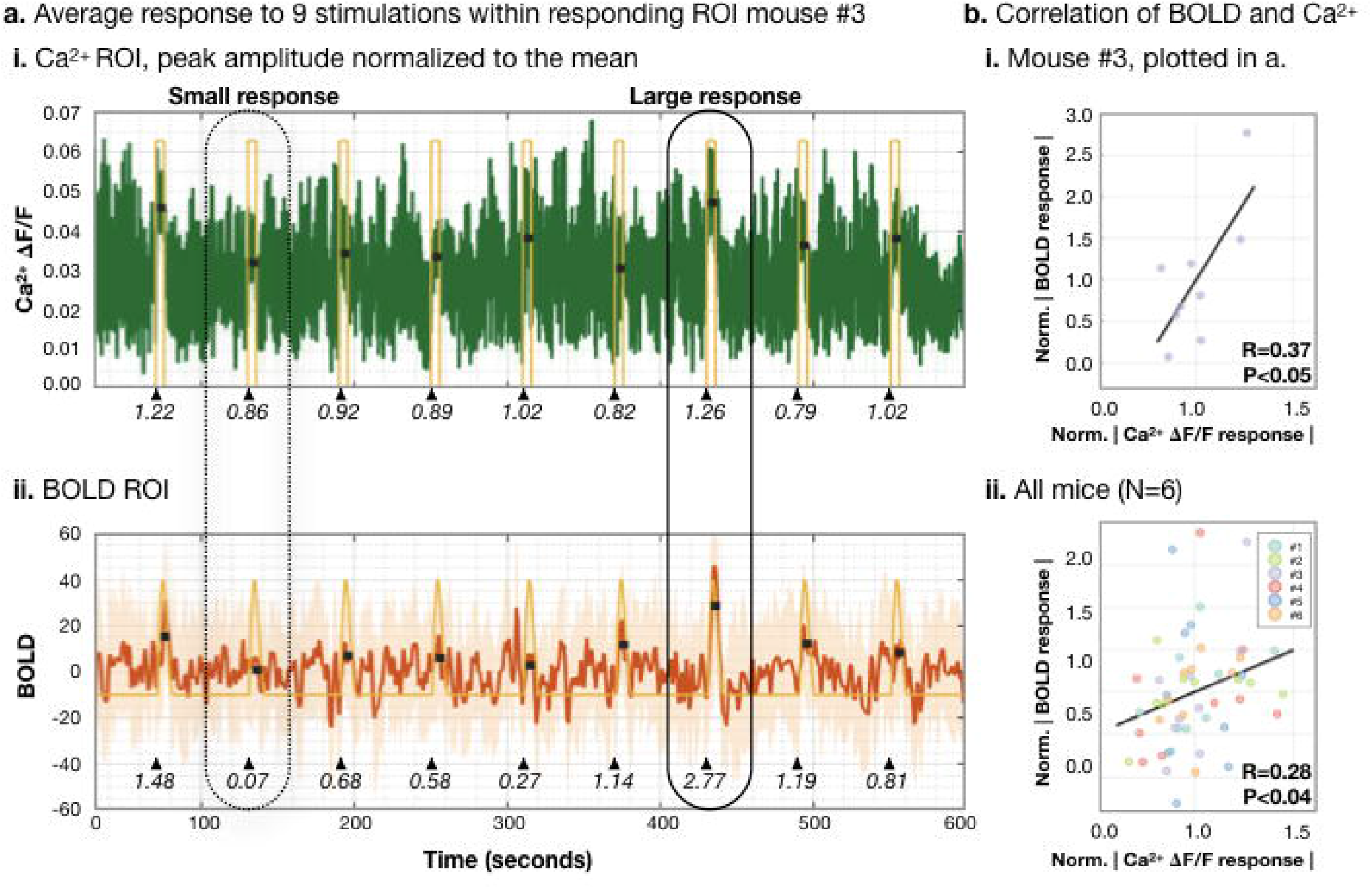
Spontaneous fluctuations of evoked Ca^2+^ and fMRI signals. The average response in the hind-paw cortex of the Ca^2+^ (green) (a.i.) and fMRI (orange) (a.ii.) data for a mouse are plotted over a 10 minute interval containing N=9 stimulation periods. For reference, the box-car (i.) and HRF (ii.) from the general linear models are plotted in yellow. During data recording there are spontaneous fluctuations in the magnitude of the Ca^2+^ and fMRI responses. Small (dashed line) and large (solid line) amplitude responses are circled and these indicate coincident fluctuations in the amplitude of the responses to the same presented stimuli. We estimate peak amplitude response by averaging the signal from the peak (see Supplementary Fig. 9 for how the peak was determined) until three seconds after the peak (each stimulus is 5 seconds long). The three second periods for each peak are shown by black lines at the level of each estimated response amplitude. The normalized response amplitude (relative to the mean signal) is written beneath each response (below black triangles which denote stimulus onset time). Notably, for this individual mouse, the normalized Ca^2+^ and fMRI response amplitudes are correlated r=0.37 P<0.05 (b.i.). Across all mice. N=6. (N=9 responses per mouse) we find the same relationship r=0.28, P<0.04 (b.ii.).

We also observed that the spatial responses of these two modalities do not completely overlap (**Supplementary Fig. 8**). This is likely a function of both the choice of statistical threshold, physiological differences between the contrasts, and differences between the modalities in spatial coverage. The BOLD fMRI signal arises from changes in local blood flow and volume in response to energy consumption that are maximal somewhat downstream of the activation area. This phenomenon has been previously observed in human fMRI studies.^35^ The measured Ca^2+^ signal is specific to excitatory neurons, whereas the fMRI response is cell-type indiscriminate and present with any energetic response, including inhibition. Furthermore, the fMRI contrast is collected in three dimensions, whereas the Ca^2+^ data is integrated across an unknown depth.

### Functional parcellation of Ca^2+^ and fMRI spontaneous activity

Here, and in remaining sections, we present results from measurements of spontaneous activity acquired in the absence of any applied stimulation. To begin, we used a data-driven parcellation procedure following a method adapted from the human fMRI literature.^36–39^ This approach uses a multi-graph k-way clustering algorithm to identify functional brain regions, referred to as nodes, which together comprise a functional atlas or parcellation (**Supplementary Fig. 10., Methods**).^40^ We applied this method using either Ca^2+^ or fMRI data to generate two functional parcellations, one per modality, for each mouse.

We performed the parcellation independently in the left and right hemispheres for both modalities. This allowed for validation of the functional parcellations by ‘folding’ across the midline and computing the similarity between hemispheres (Fig. 5 a.i.). We quantified the bilateral symmetry using the Dice coefficient (**Supplementary Material**), which showed strong interhemisphere similarity for both Ca^2+^ and fMRI data (0.68 ± 0.05 for Ca^2+^ and 0.56 ± 0.04 for fMRI data, mean ± SD, 10 parcels per hemisphere, N=6 mice) that was highly statistically significant (P<0.002) relative to randomly assigned parcel membership (Fig. 5 b). This demonstrates that both modalities show a high degree of cross-hemisphere symmetry.

**Fig. 5.**
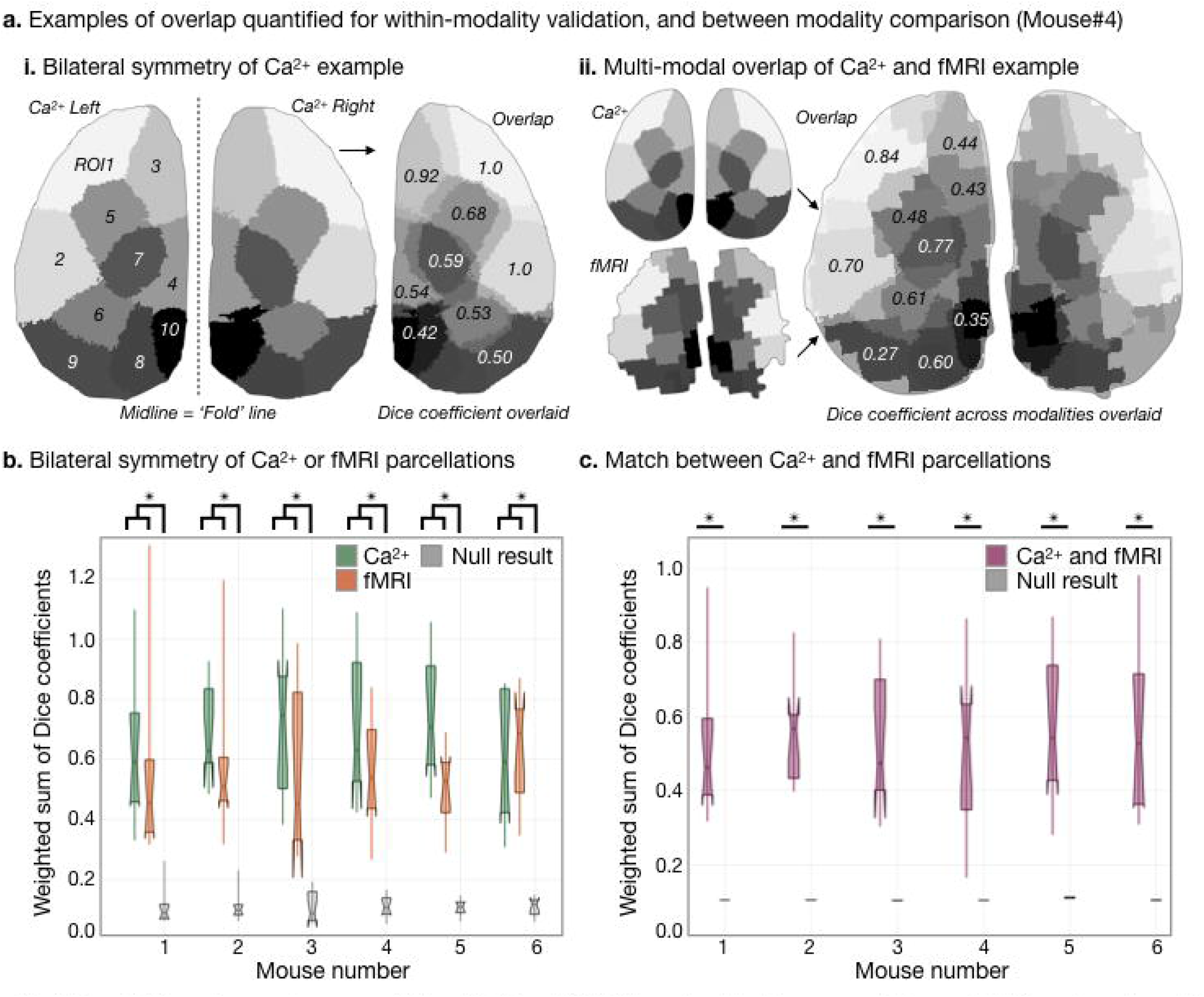
Parcellation using spontaneous activity of Ca^2+^ and fMRI. We used a data-driven approach to parcellate the cortex using multigraph k-way clustering. To measure the consistency of the functional parcellation. we separately parceled the left and right hemispheres for each modality, then compared the parcels across hemispheres (a.i.) and modalities (a.ii.) using the Dice coefficient (Supplementary Material). Bilateral symmetry of the Ca^2+^ (plotted in green), and fMRI (plotted in orange) functional parcellations was well above chance (P<0.002, null result from random parcel assignment, plotted in gray) (b.). There was also strong agreement between Ca^2+^ and fMRI parcellations (a.ii., c.), with parcels overlapping across modalities at well above chance levels for all mice (N=6, P<0.00002) (c.). While the two modalities rely on different contrast mechanisms, they are both driven by the same neural activity and thus we would expect them to yield similar results at the level of functional parcellations.

Next, we compared functional parcellations between modalities (Fig. 5 a.ii.) across the cortex, and found a strong inter-modality topological similarity that we again quantified using the Dice coefficient (0.54 ± 0.15, 10 parcels per hemisphere, N=6 mice). This relationship was also highly statistically significant (P<0.00002) relative to randomly assigned parcel membership (Fig. 5.c.), indicating that the functional parcellation of the Ca^2+^ and fMRI data identify shared brain organization. Notably, the inter-modality agreement was akin to the bilateral symmetry we observed within each modality. This result demonstrates that despite the different sources of contrast, spatiotemporal resolutions and response functions of the two modalities, the functional connectivities defined by these methods are highly consistent, as are the functional parcellations.

### Stability of Ca^2+^ and fMRI connectivity strengths over time

With functional data, and a node atlas, the correlation between the average time courses from any pair of nodes is taken as a measure of the strength of the functional connectivity between nodes. We examined the stability of Ca^2+^ and fMRI connectivity across time by calculating individualized Ca^2+^ (**Supplementary Fig. 11.a.**) and fMRI (**Supplementary Fig. 11.b.**) parcellations from five 10 minute acquisitions conducted over the course of 2.5 hours. To assess stability, we applied the parcellations to each 10 minute run separately and generated a connectivity matrix for each run, thereby yielding five connectivity profiles for each mouse, and each modality. By computing the correlation of the connectivity profiles across time (**Supplementary Fig. 11.c. and 11.d.**), we obtained a measure of connectivity profile stability during the course of an experiment. We observed that the patterns of Ca^2+^ and fMRI connectivity were very stable within each animal (Pearson correlation for Ca^2+^ r=0.995 ± 0.003, and fMRI r=0.84 ± 0.09 (mean ± SD)).

### Similarity between Ca^2+^ and fMRI connectivity strength

Parcellations based on either Ca^2+^ or fMRI data shared a similar topological pattern (Fig. 5), including correspondence to known brain regions (e.g. motor, retrosplenial, primary somatosensory, visual and auditory regions, **Supplementary Fig. 12**). To further investigate the correspondence between modalities, we transformed the Ca^2+^ parcellations onto the fMRI data, and the fMRI parcellations onto the Ca^2+^ data (Fig. 6.a.). Thus, we could examine the connectivity profiles across modalities.

**Fig. 6.**
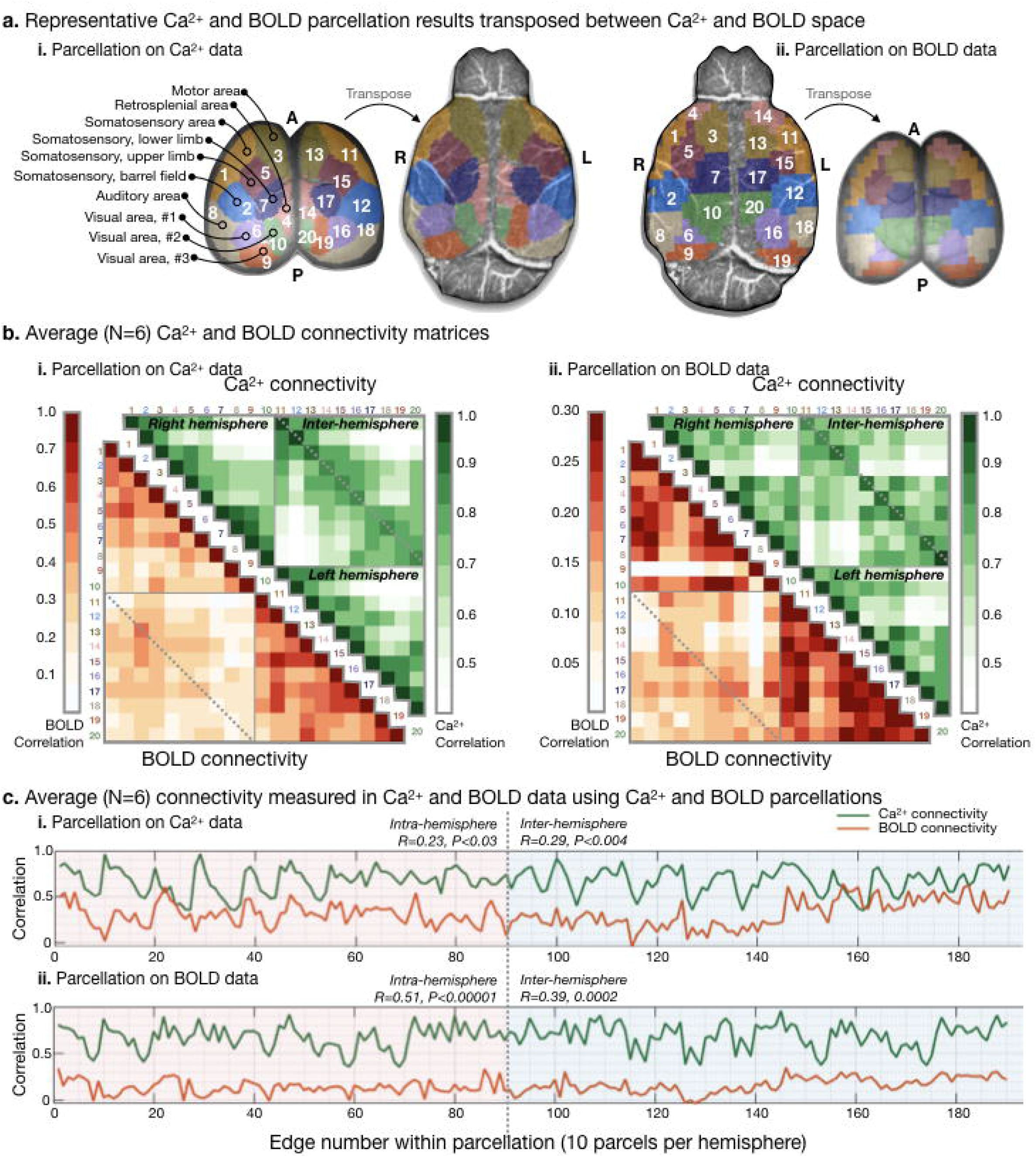
ctivity matrices for Ca^2+^ and fMRI spontaneous activity data. For all mice (N=6) functional parcellations were computed for the Ca^2+^ and fMRI data. The Ca^2+^ parcellation was projected onto the fMRI data, and vice versa. In (a.i.), we show for a single mouse, the Ca^2+^ parcellation in native Ca^2+^ space and transposed into fMRI space. Likewise, in (a.ii.), the fMRI parcellation is shown in native space as well as transposed into Ca^2+^ space. The ROIs identified within the parcellations are labeled based on proximity to regions defined by the Allen atlas (Supplementary Fig. 12). with the exception of the somatosensory lower limb area which can reliably be identified by evoked responses. In (b.)· the average (N=6) connectivity matrix for the Ca^2+^ (green, upper right) and fMRI (lower left) is shown, for the Ca^2+^ parcellation (b.i.). and for the fMRI parcellation (b.ii.). The matrices show similar patterns of strong and weak connections. In (c.). the similarity of Ca^2+^ and fMRI connectivity strength is quantified. For the Ca^2+^ (c.i.) and fMRI (c.ii.) parcellations. the average Ca^2+^ (green) and fMRI (orange) connectivity strength is plotted. Intra-hemisphere (shaded in pink), inter-hemisphere (shaded in blue), and whole brain Ca^2+^ and fMRI connectivity are correlated. Thus, regions identified by either Ca^2+^ or fMRI based parcellation which show high/low synchrony in Ca^2+^ also show the same patterns in fMRI. Functional connectivity as measured by fMRI has not previously been validated by an independent imaging modality such as this Ca^2+^ mesoscopic imaging approach.

From each parcellation, we computed a connectivity matrix for each data set using each (Ca^2+^ or fMRI) data-driven parcellation (yielded four connectivity matrices) (Fig. 6.b.). Regardless of whether the parcels were calculated from Ca^2+^ or fMRI data, the within hemisphere fMRI connectivity strengths were greater (mean r=0.20) than the between hemisphere connectivity strength (mean r=0.10; P<0.05), whereas there was no difference in the within versus between hemisphere Ca^2+^ connectivity strength (mean r=0.67; P>0.1). This finding is similar to previous observations using voltage indicators and optical measurements of the hemodynamic signal (a proxy for BOLD), where the bilateral symmetry of the voltage indicators was found to be greater than the bilateral symmetry of the hemodynamic signal.^41^ However, we also observed that bilaterally paired regions (e.g. primary somatosensory areas in the right and left hemispheres) were more correlated than mismatched regions (e.g. primary somatosensory and visual areas), for both modalities Fig. 6.b.

To quantify the similarity between Ca^2+^ and fMRI spontaneous activity patterns across the cortex, we computed the correlation between Ca^2+^ and fMRI connectivity profiles (Fig. 6.c.). For both Ca^2+^ and fMRI parcellations, inter-/intra-hemisphere and whole brain connectivity showed a relationship between modalities. Regions which showed high/low connection strength in Ca^2+^ data also showed high/low connection strength in fMRI data (Fig. 6.c.), P<0.03. This result demonstrates that the Ca^2+^ and fMRI signals are providing partially shared information, and gives strong evidence of a neural basis for the functional connectivity metrics obtained from resting-state fMRI data. Indeed, this result is a unique affirmation that both the organization which we can derive from spontaneous fMRI activity, and the functional relationships between the regions, have neurological underpinnings. Until now, this has been a long-held assumption with no validation.

## DISCUSSION

Functional connectivity as measured by fMRI was first introduced in 1995.^42^ Ever since, the field has struggled to determine whether these signals reflect meaningful functional connections or simply some form of correlated noise. Converging evidence suggests that these temporal correlations do reflect neural connections but, despite its wide-spread use, functional connectivity as measured by fMRI has been extremely difficult to independently validate. This challenge stems from the technological difficulty of implementing an independent imaging approach that can be run simultaneously with fMRI. The multimodal approach presented here provides such a methodology, and both the Ca^2+^ and fMR imaging reveal functional connections spanning the cortex with very similar connectivity matrices. This demonstrates that these independent indices of correlated activity share an underlying neural signal source.

No single imaging modality can provide the requisite information needed to connect cellular activity to brain-wide network activity. Serial experiments using different modalities, even in the same model, are insufficient to unequivocally establish links between different measurements. This is due to the complexity of the brain, and the difficulty of controlling all the factors that influence activity. A method to circumvent this challenge has been to align evoked responses across experiments. As we demonstrated here, however, there are biologically meaningful differences between individual responses to identical stimuli. Our finding that the Ca^2+^ and fMRI evoked response amplitudes are correlated dispels the notion that response-to-response fluctuations are caused solely by experimental noise. Rather, subtle differences in both signals, which are typically removed by averaging, are in fact measurements of true moment-to-moment changes in brain activity.

Evoked responses provide data for determining the spatiotemporal transfer functions between modalities. While Ca^2+^ has much higher spatiotemporal resolution and provides potentially better localization of neural activity, the evoked responses of the two modalities have similar response functions. The strength of the Ca^2+^ signals in this context is in the spatiotemporal resolution as well as the selective specificity to different cell types, whereas the strength of the fMRI lies in whole-brain coverage (including structures deep within the brain), while also providing a link to human neuroscience research. In future experiments, measuring evoked responses to different sensory stimuli will allow the calculation of cell-type specific transfer functions between modalities.^43,44^ In addition, elucidating these relationships will provide a means for matching cortical activity to activity within deeper brain regions.^45,46^

Limiting studies to only evoked responses is problematic because it ignores the vast majority of brain activity that is spontaneous or independent of applied stimulation. Spontaneous activity cannot be studied across spatiotemporal scales in separate experiments without a means for linking activity between scales. This challenge motivates the need for multi-modal methods with wide spatiotemporal ranges, which are essential to gaining a comprehensive understanding of the relationships that govern brain function across scales.

Fluorescence imaging using single fibers in combination with fMRI, or wide-field fluorescence and intrinsic hemodynamic signal imaging, have been described previously.^23–28,47–49^ Both these dual imaging approaches measure Ca^2+^ and BOLD (or a proxy for BOLD), but lack spatial coverage. Single-fiber time-series data, while quite useful for studying evoked responses, cannot reveal large scale network interactions compared to the whole-cortex Ca^2+^ imaging presented here. Likewise, optical imaging methods struggle to access regions beneath the upper cortical layers, while fMRI provides the ability to image throughout the depth of the brain and in three dimensions. The method presented here is uniquely able to simultaneously measure both the intrinsic hemodynamic signal and fMRI at high resolution and an extended spatial scale.

This new methodology measures concurrent activity from two independent sources both related to aspects of neural activity. Using spontaneous activity measurements, we generated functional parcellations for both the Ca^2+^ and fMRI data. The resultant connectivity matrices revealed similar topographic structure that was highly stable across the duration of our study. These results reveal correspondence between the fundamental structural and temporal patterns of these two independent signal sources arising from spontaneous activity. This represents the first independent imaging validation of the neural basis for functional connectivity as measured via fMRI and affirms the long-held assumption about the neural contributions to this signal. Given that the Ca^2+^ and fMRI signals arising from both evoked and spontaneous activity fundamentally measure different correlates of brain activity, we did not expect these measurements to be identical. Indeed, perfect correspondence would abrogate the need to perform dual-modality simultaneous imaging.

A salient feature of our results is that at the level of mapping cortical networks using spontaneous signals there is good agreement between nodes derived from fMRI and Ca^2+^ signals, whereas for sensory-evoked responses the overlap between foci derived from fMRI and Ca^2+^ signals was more marginal. This apparent discrepancy indicates that the point-spread function (PSF) of each modality differs owing to the distinct origins of fMRI and Ca^2+^ signals.

Fundamentally, the fMRI signal arises from a complex combination of functional hyperemic events, whereas the Ca^2+^ signal represents neuronal-specific events that are dominated by signals originating from the neuropil. While we can safely exclude confounding contributions of large blood vessels for BOLD contrast at 11.7T, the PSF for fMRI is still slightly larger than the nominal voxel size because of the echo-planar imaging used in our experiments.^50^ Hence the PSF for fMRI is spatially smoothed beyond the 3D isotropic voxel. Although the in-plane spatiotemporal resolution of the Ca^2+^ imaging data is significantly higher than fMRI, the PSF for Ca^2+^ imaging is complex. The 2D voxel has an unknown contribution in the depth dimension, and it is also likely that the intensity in each 2D voxel may have contributions from memory effects due to the high temporal resolution imaging. Future experiments will be designed to specifically resolve the differing spatiotemporal properties of the fMRI and Ca^2+^ signals.

While only excitatory neurons were examined in the Ca^2+^ imaging experiments presented here, a key strength of this dual imaging approach is the capability of measuring the relative contributions of different cell populations to the fMRI signal. Little is known about cell-specific contributions underlying the macroscopic functional organization of brain function. Similarly, the individual contributions of specific cell populations to the fMRI signal has never been studied, but is now possible with this methodology. Emerging technologies such as voltage sensitive indicators could be used instead of Ca^2+^ sensors, providing further insight into the link between cell level function and meso- or macroscopic network level brain organization.^51,52^ This methodology will provide substantial mechanistic insight into the feed-forward and feed-back communication between brain-wide functional motifs and cellular activity, and there is tremendous value in obtaining multi-modal data from mice expressing fluorescence indicators within different cell populations and applying this methodology to imaging with voltage indicators.

Our initial experiments were performed under light isoflurane anesthesia because of the additional challenges involved in imaging awake animals in the MR environment. Even light anesthesia can have profound effects on brain activity, but the anesthetized model was sufficient for demonstrating the correspondence between Ca^2+^ and fMRI signals and proof-of-principle for this new dual-modality imaging tool.^53,54^ All features of the device and procedures described here are compatible with imaging awake animals. Furthermore, the methods described are also suitable for longitudinal imaging studies.

Given that mice were anaesthetized, it was unclear if the connectivity profiles would be stable. Nevertheless, the functional connectivity patterns that emerged were highly stable despite the long duration of the experiments and the challenge of maintaining the physiological stability of small animals within the MR scanner. The stability of the multi-modal functional metrics observed here in lightly anesthetized animals is encouraging as we move forward with developing awake imaging protocols to investigate state-dependent changes in functional connectivity.

Overall, the novel hardware and software developments presented here have broad and generalizable applicability. In addition to the research avenues outlined above, our simultaneous measurements can be used to validate the application of graph theory approaches in the analysis of fMRI data – a rapidly emerging area of research in human fMRI. Specifically, these data can be used to test approaches that aim to extract dynamic connectivity components using the optical signal as a ground truth measure.^55–57^ In human fMRI research, these endeavors have received much attention.^36,58–65^ Thus, data obtained using our approach will impact a wide audience of clinical and basic science researchers. Finally, our method can be applied in neurological models of disease or injury to measure pathology and to test intervention strategies.

The present work is the culmination of a highly innovative and interdisciplinary effort to span spatiotemporal scales from the micro-to macroscopic-scale.^66^ Fundamentally, this simultaneous imaging method will provide data that contributes to a firmer biological understanding of the fundamental cellular origins of many of the macroscopic signal changes observed with fMRI. The fMRI component of this multimodal imaging approach provides a direct link from mouse to human studies, giving fundamental insights into the functional organizing principles of the brain across species in both health and disease as well as throughout the lifespan.

## METHODS

### Surgical preparation

All procedures were performed in accordance with the Yale Institutional Animal Care and Use Committee and are in agreement with the National Institute of Health Guide for the Care and Use of Laboratory Animals. All surgical materials are compatible with MRI. Anesthesia is induced with 3% Isoflurane (1.5-2% during surgery). Fur is removed from the scalp and thigh (for MouseOx) using dilapidation cream (Nair™). Lidocaine (0.5%, Henry Schein Animal Health VINB-0024-6800) and marcaine/epinephrine (0.5%, Pfizer Injectables 00409175550) are used to numb the scalp prior to resection. Once the skull surface has been cleared of tissue, a dental cement (C&B Metabond^®^, Parkell) ‘well’ is built around the circumference of the skull surface; taking care not obstruct the Ca^2+^ imaging FOV. A fluorescent bead (Fluorescent green PE microspheres, UVMS-BG-1.00, 106-125μm, Cospheric) is embedded within the right-anterior wall for motion correction. Dental cement is also used to secure the outside edges of the well to our in-house built head-plate (acrylonitrile butadiene styrene plastic, Lulzbot TAZ-5 printer, with 0.35mm nozzle) which cradles the sides of the skull and attaches above the olfactory bulb.

The head-plate dovetails with the RF MRI coil and Ca^2+^ imaging hardware to minimize motion and aid alignment. The well is filled with an optically transparent agar substitute: 0.5% Phytagel (BioReagent, CAS.71010-52-1) with 0.5% MgSO4 in water, sealed with a glass cover-slip (Carolina Biological Supply Company, item.no.633029) and secured with dental cement. The well is necessary to avoid artifacts in the MRI data which are caused by neighboring materials with large differences in magnetic susceptibility (eg. a skull to air interface). Furthermore, the glass cover-slip provides a smooth surface which improves Ca^2+^ signal transmission. Care must be taken during the preparation to eliminate bubbles in the dental cement and Phytagel which cause artifacts in the MRI and Ca^2+^ data.

### Optical components

The optical components of the imaging system were composed of glass or plastic (**Supplementary Fig. 2**). Directly above the mouse, a prism (25mm, uncoated, N-BK7, #32-336 RA-Prism, Edmund Optics) redirected the excitation (entering) and emission (exiting) light by 90° so that they pass through the telecentric lens (MML-1-HR65DVI-5M, Moritex), which houses several optical components. For MRI compatibility, we replaced the metal housing of the stock telecentric lens with plastic components (Derlin^®^ Acetal Resin and PEEK, McMater-Carr). We further customized the telecentric lens by replacing the beam-splitter with a dichroic filter (15×17×1mm, 495nm high-pass, T495lpxr, Lot: 321390, CHROMA) in order to appropriately direct the excitation and emission light. A custom port that collimates (12 Dia. x 15 FL mm, VIS-EXT, Inked, Plano-Convex, Edmund Optics) and redirects the excitation light by 90° into the telecentric lens was purpose built out of Delrin and nylon (Kramer Scientific). The excitation light arrives via a 5mm x 5m liquid light guide (10-10645, Lumencor) from the room neighboring the magnet where the LED light source (Lumencor SPECTRA X, Lumencor) is housed. Similarly, a custom 4.6m long 14.5 × 14.5mm^2^ cross section coherent fiber optic bundle (N.A. 0.64) containing an array of multifibers (10μm elements in a 6×6, 60μm X 60μm array for a total of nearly 2,000,000 fibers) (SCHOTT Inc.) transports the emission light to a room neighboring the magnet where the Ca^2+^ imaging data is recorded using a sCMOS camera (512×512 pixels, pco.edge 4.2, PCO). An additional GFP emission filter (ET525/50m, CHROMA) is placed between the optical fiber bundle and the camera connected by two optical extenders (sub-assemblies of a TwinCam LS Image splitter, Cairn Research).

### Mice

Mice were housed on a 12 hour light/dark cycle. Food and water were available ad libitum. Mice were adults, 6-8 weeks old, 25-30g, at the time of imaging. We report data from a first-generation TIGRE (genomic locus) line crossed with reporter lines controlled by Cre recombinase (promoter) with a tetracycline-regulated transcriptional trans-activator (tTA) for amplification.^19–22^ For detailed expression level data, refer to: http://connectivity.brain-map.org/transgenic;http://www.alleninstitute.org/what-we-do/brainscience/research/products-tools/.^19^ More specifically, we used Ai93 mice (or TIT2L-GCaMP6f, <u>T</u>IGRE - <u>I</u>nsulators - <u>T</u>RE2 promoter - <u>L</u>oxPStop1LoxP - <u>G</u>FP <u>C</u>al<u>M</u>odulin fusion <u>P</u>rotein 6 fast) crossed to CaMK2a-tTA mice (<u>C</u>al<u>M</u>odulin dependent protein <u>K</u>inase <u>2</u> <u>a</u>lpha) driven by Slc17a7 (or Vglut1) - IRES (<u>i</u>nternal <u>r</u>ibosome <u>e</u>ntry <u>s</u>ite) 2 - Cre promotor mice, all purchased from Jackson Labs (JAX stock numbers: 024103, 003010 and 023517) and bred in-house. The resulting Slc17a7-Cre; Camk2a-tTA; Ai93 mice have GCaMP6f expressed in cortical excitatory neurons.

### Animal monitoring

During data acquisition, mice (N=6) were minimally anesthetized with 0.5-1.25% Isoflurane, adjusted to maintain a heart rate of 480-550 beats per minute. Mice freely breath a mixture of O2 and medical air, adjusted to maintain an arterial O2 saturation of 94-98%. Heart and breath rate, arterial O2 saturation, and rectal temperature, were continually monitored (MouseOx from STARRLife Sciences, Inc.) and recorded (Spike2, Cambridge Electronic Design Limited) (Supplementary Fig. 3.). Body temperature was maintained with a circulating water bath. During image acquisition, MRI, Ca^2+^ imaging and physiological data recording were synchronized (Master-8 A.M.P.I., Spike2 Cambridge Electronic Design Limited). Hind-paw electrical stimulation was delivered at 1mA, 5Hz, in 5/55 seconds ON/OFF cycles.

### Functional imaging parameters

**Ca^2+^ data** is recorded at an effective rate of 10Hz. To enable frame-by-frame background correction (next section), violet (395/25) and cyan (470/24) illumination is interleaved at a rate of 20Hz. The exposure time for each channel (violet/cyan) is 40ms to avoid artifacts caused by the rolling shutter refreshing. Thus, the sequence is as follows: 10ms blank, 40ms violet, 10ms blank, 40ms cyan etc.

**fMRI data** is acquired using a gradient-echo, echo-planar-imaging (EPI) sequence with a repetition time (TR) of 1 second, and an echo time (TE) of 9ms. The data are collected at a 0.4×0.4×0.4mm^3^ resolution, across 28 slices; yielding whole brain coverage. Each functional imaging run is 600 repetitions in length (10 minutes).

### Ca^2+^ data processing

Images are rotated to align the anterior-posterior axis with vertical (MATLAB, *imrotate*). Violet and cyan frames, which are interleaved during data acquisition, are separated (odd/violet and even/cyan) and processed in parallel. Motion correction, using rigid body translation (MATLAB, *imregtform*), is performed on even/odd frames using a FOV containing the fluorescent bead (not brain tissue) imbedded within the dental cement (Fluorescent green PE microspheres, UVMS-BG-1.00, 106-125μm, Cospheric; **Supplementary Fig. 4**).

Data are masked to isolate brain tissue (ImageJ, *polygon*). For even/odd frames, individual pixel baseline correction (MATLAB *imtophat*, line structure 300 width) is applied to remove the trend caused by photobleaching during the experiment. This results in a zero baseline. Data are baseline shifted back to the raw-data signal intensity by adding the average (pre-baseline correction) signal of each pixel’s time course back to each pixel value. This is necessary for regression of the background signal (next step) and calculating the relative fluorescence change (final step). To remove non-GCaMP6f fluorescence changes (e.g. hemodynamic signal), we do background correction by regressing (MATLAB, *regress*) the violet from the cyan time course pixel-wise.^29,30^ To compute the relative fluorescence change, each pixel in the background corrected (violet regressed from cyan) time course is divided by the average signal of each pixel’s time course. The result is the ΔF/F (F, fluorescence) movie (**Supplementary Fig. 4**).

### MR surface projection image (2D)

The MR surface projection image, is generated using a custom ray-casting algorithm on the masked angiography image (Fig. 2.c.i.) to create a 2D projection image (Fig. 2.c.ii.). This algorithm projects rays from the top of the image and uses the brain surface normal for shading to create a synthetic brain surface image (**Supplementary Fig. 6., Supplementary Material**). This effectively projects the MR into the same space as the Ca^2+^ data which is viewed from above. This image contains anatomical details (e.g. cortical surface vessels) that are also visible in the average Ca^2+^ image. For each mouse, anatomical features present in both images, such as the projections of the middle cerebral arteries and the midline, are used to generate a rigid transformation that brings the Ca^2+^ and MRI data into alignment (Fig. 2.c.iv.).

### fMRI data processing

Data are motion corrected (AFNI, Analysis of Functional NeuroImages, *3dVolReg*), masked to isolate brain tissue (MATLAB, *roipoly*), and spatially blurred within the brain-mask (MATLAB, *smooth-gaussian*, full-width-half-maximum, FWHM, 0.8mm).^31^ The data are filtered (0.01-0.2Hz, MATLAB, *butterworth*), the global signal regressed (MATLAB, *detrend*), and the linear trend removed (MATLAB, *detrend*). Data with frame-wise motion estimates >0.4mm are excluded (voxel size 0.4×0.4×0.4mm^3^). The majority of the data we collect contains sub-threshold motion (75% of evoked activity recordings and 82% spontaneous activity recordings, N=28/38), Supplementary Fig. 5.

### Structural MR-imaging parameters

**High in-plane resolution images of fMRI FOV**. Using a multi-spin-multi-echo (MSME) imaging sequence. In 10 minutes, 40 seconds, using a TR/TE of 2500/20ms, we obtain 28 slices (0.4mm thick) with an in-plane resolution of 0.1×0.1mm^2^ (two averages). The slice prescription of these images matches those of the functional MR-images (i.e. they are of the same anatomy).

**Isotropic 3D anatomy of the whole brain**. Using a MSME imaging sequence. In 5 minutes, 20 seconds, using a TR/TE of 5500/15ms, we obtain a 0.2×0.2×0.2mm^3^ (single average) image of the whole brain. This sequence is repeated five times during our imaging protocol interleaved with functional acquisitions. Interleaving structural and functional acquisitions allows recovery of the Ca^2+^ signal and more robust responses to stimulation during functional data acquisitions with evoked responses. In post-processing, the five isotropic anatomical images are concatenated (MATLAB, *horzcat*), motion correct (AFNI, *3dVolReg*), and averaged (MATLAB, *mean*) to create one image.

***MR-angiogram***. Using a fast-low-angle-shot (FLASH) time-of-flight (TOF) imaging sequence. In 18 minutes, using a TR/TE of 130/4ms, we obtain a 0.05×0.05×0.05mm^3^ 2.0×1.0×2.5cm^3^ image of the blood vessels within the cortex.

**High resolution anatomy of angiogram FOV**. Also using a FLASH sequence. In 7 minutes, 30 seconds, using a TR/TE of 61/7.5ms, we obtain an image with resolution 0.13×0.08×0.05mm^3^ and FOV 2.0×1.0×2.5cm^3^ capturing the anatomy within the same FOV as the MR-angiogram.

### Generalized linear model (GLM) for Ca^2+^ and fMRI

**Ca^2+^ data.** From each mouse, we collect 40 minutes of evoked responses to hind-paw stimulation during four 10 minute sessions which are interspersed between spontaneous activity recordings and structural imaging. Preprocessed Ca^2+^ data containing evoked responses are normalized (MALTAB, *zscore*) and fit using a GLM (MATLAB, *glmfit*). Motion parameter estimates from simultaneously recorded fMRI data, a drift parameter (photobleaching), motion estimates from fluorescent beads (**Supplementary Fig. 4**), as well as a box-car response function are included in the model. The response map is thresholded for pixels with beta values larger than the FWHM beta value. Clusters with <30 pixels are discarded (MATLAB, *bwconncomp*). Similar ROI results are obtained when motion estimates and/or drift parameters are not included within the model.

**Functional MRI data.** Data containing evoked responses are fit using a GLM (AFNI, *3dDeconvolve*). Drift and motion parameters as well as a custom hemodynamic response function (HRF) are included in the model. The HRF is derived from the average evoked response recorded from a mouse imaged during a pilot experiment (not one of the N=6 reported in Results), refer to **Supplementary Material** for the derivation of the HRF. The response map is thresholded to correct for multiple comparisons (false discovery rate q<0.01) and a cluster size limit (>30 contiguous voxels) applied.

### Ca^2+^ and fMRI parcellation: multi-graph k-way clustering

We denote the original data set (either Ca^2+^ or fMRI) as 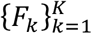, where k can be the index for different runs from one animal or the index for different animals. Each *F_k_* is organized as a 2D matrix *F_k_* = [*f*_1*k*_*f*_2*k*_ … *f_ik_* … *f_Nk_*], where each column is a time course indexed by i, and N is the total number of pixels (2D, Ca^2+^ data) or voxels (3D, fMRI data).

To apply the clustering algorithm, we construct a graph 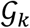 for each set of data in *F_k_*. The graph consists of vertices and edges. Vertices are the N pixels/voxels and edges are the connections between each pair of vertices. Edges are characterized by their strength, which is quantified by measuring the similarity between the time courses of pairs of vertices. Accordingly, we calculate a matrix of weights *W_k_* of size N x N for a given *F_k_*, and each entry *w_ij_* is defined by 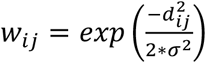. Here, we define 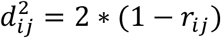, where *r_ij_* is the Pearson correlation between the time course of pixel/voxel i and pixel/voxel j.

The optimization and computation of the clustering algorithm is performed in the spectral domain. In other words, given a *W_k_*, we compute the first m eigenvectors of *W_k_* denoted as *X_k_* = [*x*_1*k*_*x*_2*k*_ … *x_mk_*], *X_k_* is of size N x m, and each column is an eigenvector. The multi-graph k-way clustering algorithm is then set to solve the following optimization,

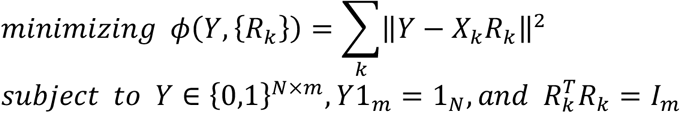

where Y is the m-ROI parcellation of the brain based on the complete sets of data from all K runs, 1. is a single column vector of size N, and *I_m_* is the m x m identity matrix. The optimization is solved iteratively. For more details refer to Shen et al. 2013.^40^ After Y is solved, we convert Y to a 1D label. Each row of Y corresponds to one pixel/voxel. By definition, each row of Y has one out of m entries equal to one, and all other (m-1) entries are zero. Thus, the label for each pixel/voxel i is the column index where the i^th^ row of Y equals one. Finally, the 1D label is mapped to the original 2D/3D space for visualization and further analysis, **Supplementary Fig. 10.**

