## Supplementary Fig. 1. for "Simultaneous mesoscopic Ca^2+^ imaging and fMRI: Neuroimaging spanning spatiotemporal scales"

### **Supplementary Fig. 1.** Assembly of MR-saddle coil, mouse head-plate, and $Ca^{2+}$ imaging optical apparatus

**a.** Removable saddle coil, with case for hardware

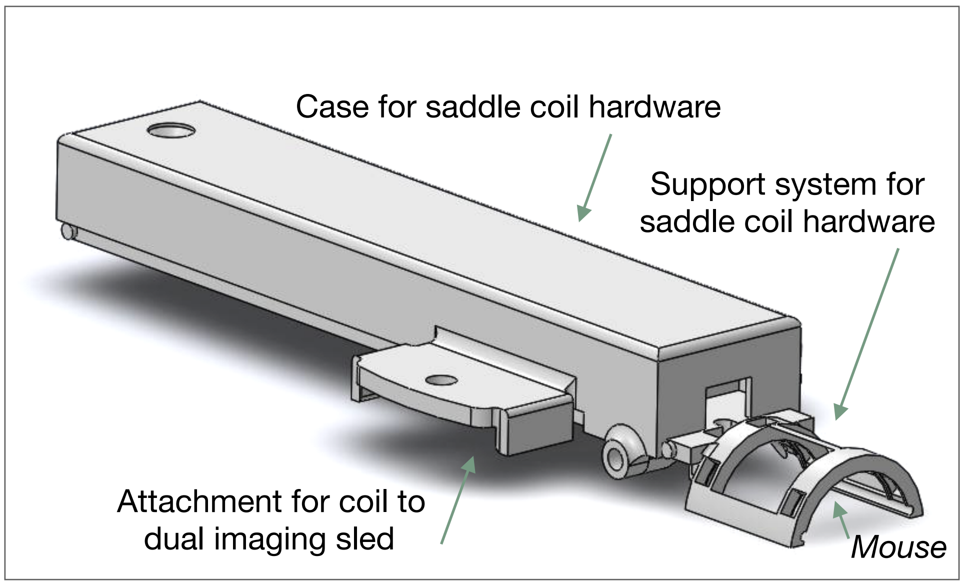


**b.** Coil in place on dual imaging sled

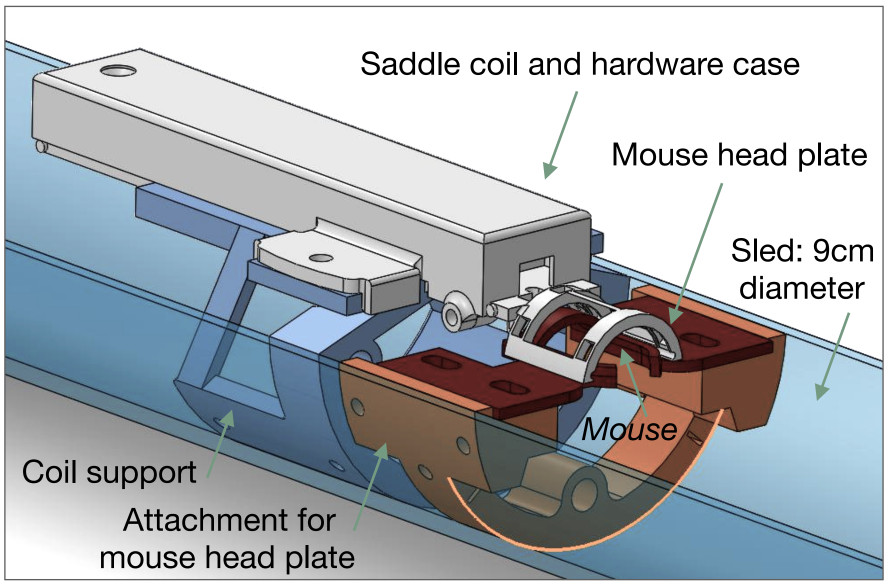


**c.** Assembled dual imaging apparatus

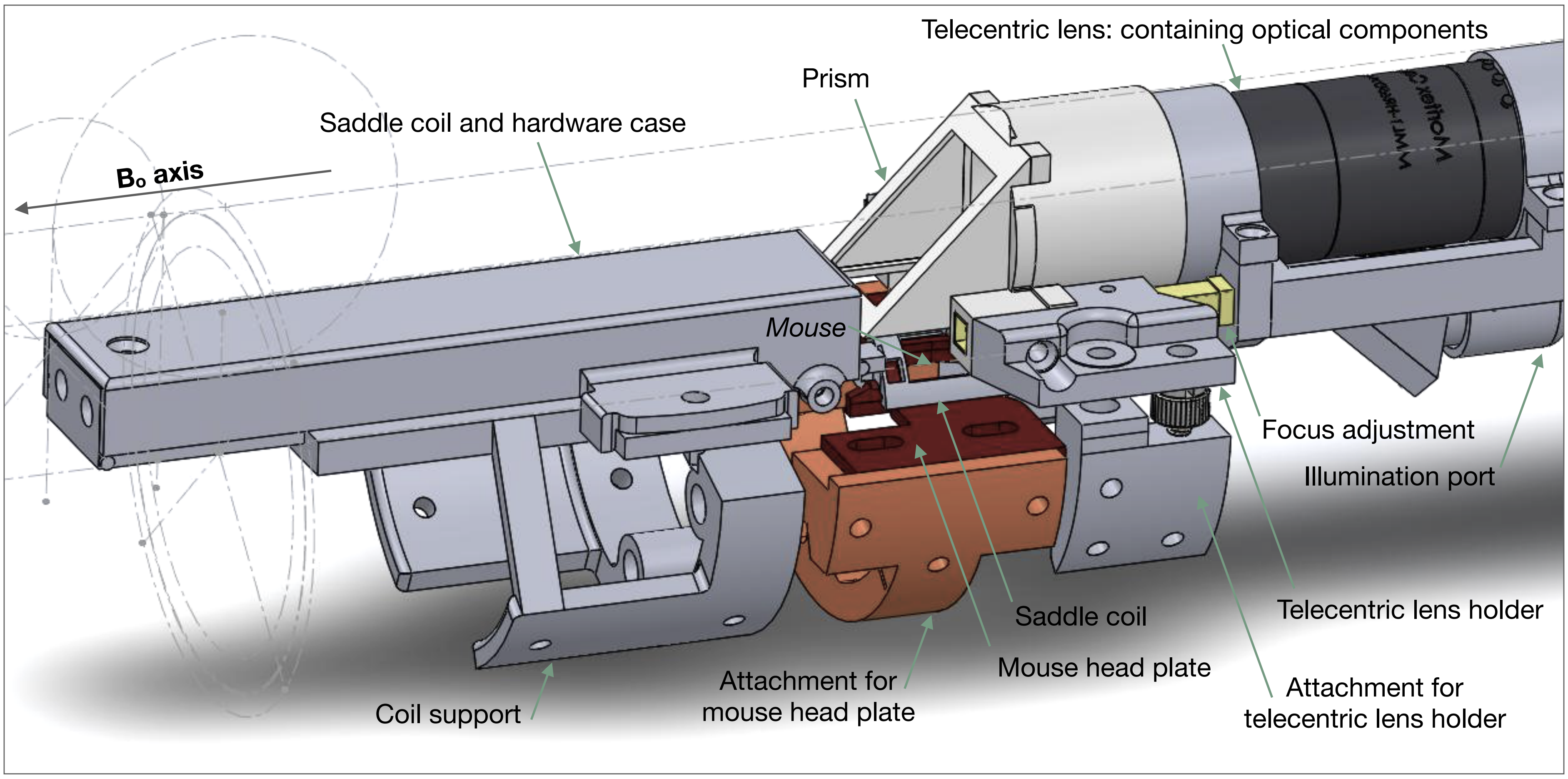


**Supplementary Fig. 1. Assembly of MR-saddle coil, mouse head-plate and $Ca^{2+}$ imaging optical apparatus.** For simultaneous imaging, we built a custom saddle coil for MR-signal reception (a.). The saddle coil is well suited to the simultaneous imaging experiment as it does not obstruct the $Ca^{2+}$ FOV, has uniform MR-signal sensitivity across the whole brain, and is relatively easy to construct. The coil case and hardware are removable which eases the placement of the animal, and allows customization of the coil for different animal sizes. (b.) The coil is mounted on a support system (blue) which is fixed to the sled. Similarly, there is a support system (orange) attached to the sled to which the mouse head plate (red) attaches. (c.) Finally, the telecentric lens (for $Ca^{2+}$ imaging) is secured above the mouse and saddle coil. The position of the telecentric lens (and housing) can be adjusted (yellow) along the magnet $B_o$ axis to focus the $Ca^{2+}$ image.
