## Supplementary Fig. 2. for "Simultaneous mesoscopic Ca^2+^ imaging and fMRI: Neuroimaging spanning spatiotemporal scales"

**Supplementary Fig. 2.** A cross section of the optical apparatus

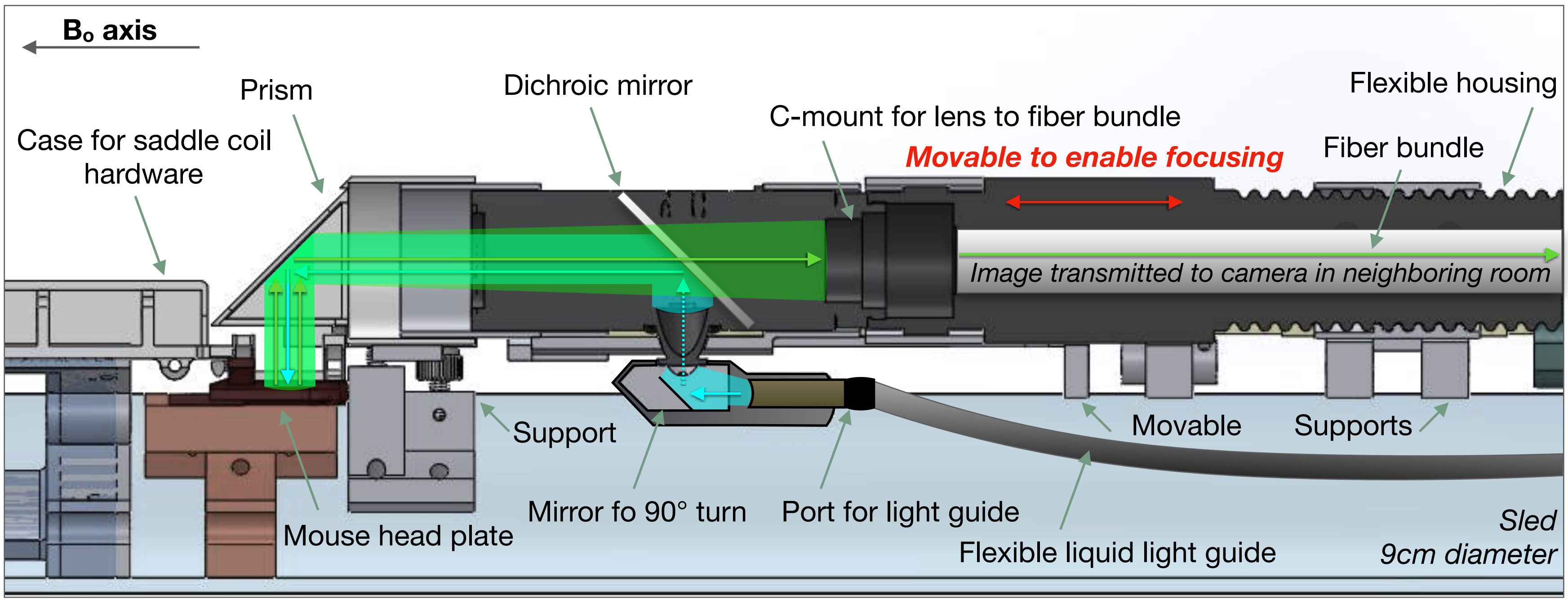


**Supplementary Fig. 2. A cross section of the optical apparatus.** To collect $Ca^{2+}$ data within the scanner necessitates redirecting both the excitation and emission light so that it can travel along the 9cm bore of the magnet. A diagram of the light path overlaid on a cross section of our optical apparatus is shown. The light enters the system via a flexible liquid light guide. At the base of the telecentric lens, the light is bent by 90° degrees to enter the telecentric lens. Upon entering the telecentric lens, the excitation light reflects off of the dichroic mirror and is redirected along the length of the telecentric lens and into the prism at the end of the apparatus. The prism redirects the excitation light onto the mouse cortex. The emission light is similarly re-directed by the prism along the length of the telecentric lens (this time traveling in the opposite direction), and passes through the dichroic mirror. The fiber bundle array is mounted onto the end of the telecentric lens and transmits the light to the room neighboring the magnet where the camera is housed. Importantly, we are able to move the fiber bundle relative to the telecentric lens to focus our recordings (red arrows).
