## Supplementary Fig. 3. for "Simultaneous mesoscopic Ca^2+^ imaging and fMRI: Neuroimaging spanning spatiotemporal scales"

### **Supplementary Fig. 3.** Physiological monitoring during simultaneous mesoscopic $Ca^{2+}$ and MR-imaging

**a.** Mean and standard deviation of physiological parameters recorded during dual imaging experiments

| | Heart Rate (bpm) | Breath Rate (bpm) | Arterial $O_2$ saturation (%) | Rectal temperature (°C) |
|---|---|---|---|---|
| **Mouse #1** | 470 ± 8 | 69 ± 20 | 98.3 ± 0.3 | 37.8 ± 0.1 |
| **Mouse #2** | 512 ± 4 | 136 ± 19 | 96.5 ± 0.8 | 38.1 ± 0.1 |
| **Mouse #3** | 487 ± 9 | 76 ± 28 | 97.9 ± 0.3 | 38.1 ± 0.2 |
| **Mouse #4** | 481 ± 15 | 102 ± 36 | 95.7 ± 1.0 | 38.1 ± 0.1 |
| **Mouse #5** | 528 ± 15 | 102 ± 15 | 98.0 ± 0.4 | 37.6 ± 0.2 |
| **Mouse #6** | 503 ± 70 | 120 ± 65 | 98.9 ± 0.4 | 37.6 ± 0.4 |

**b.** Sample physiological monitoring results during one 10 minute acquisition

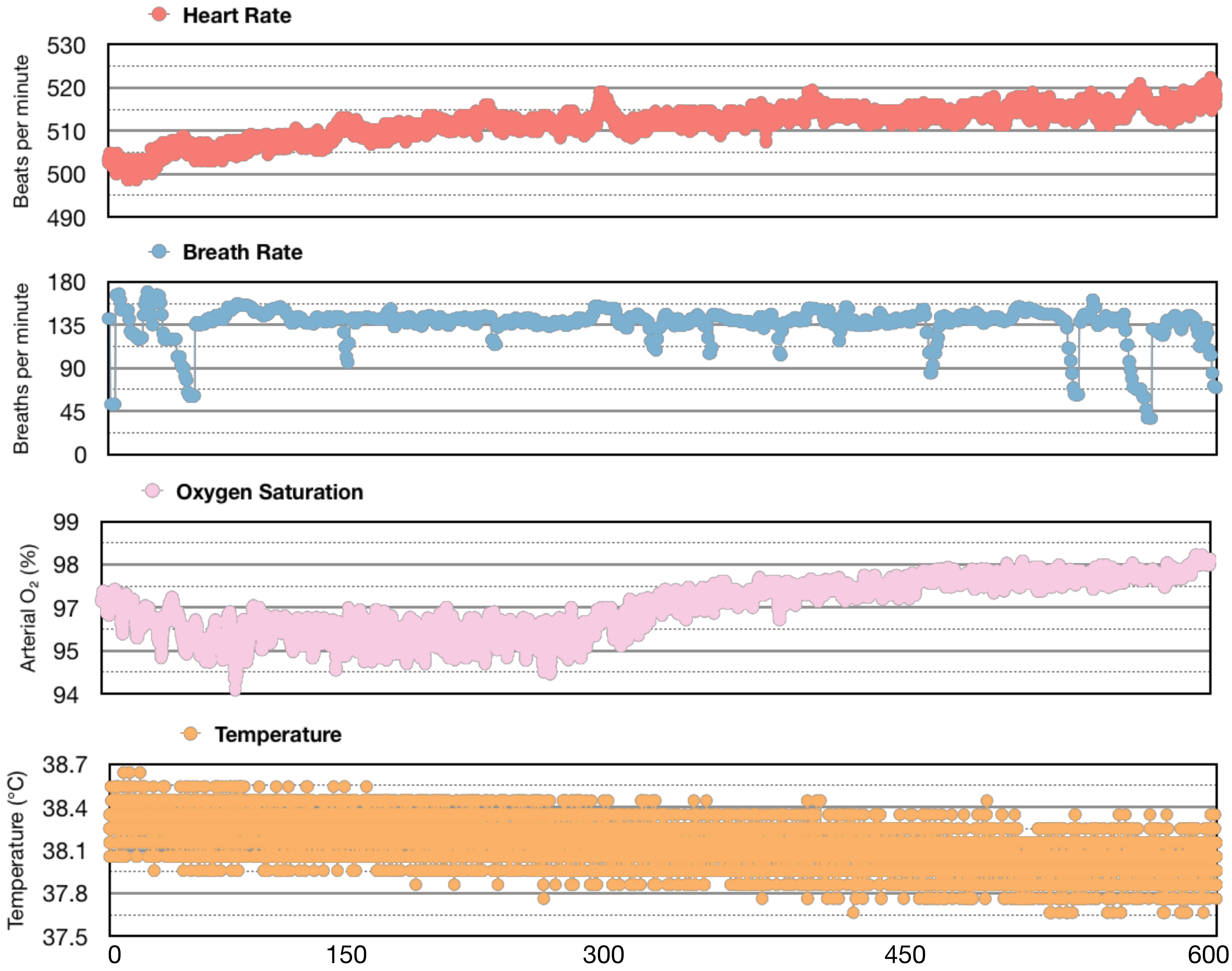


**c.** Mean MR signal across the whole brain (data is regressed during pre-processing)

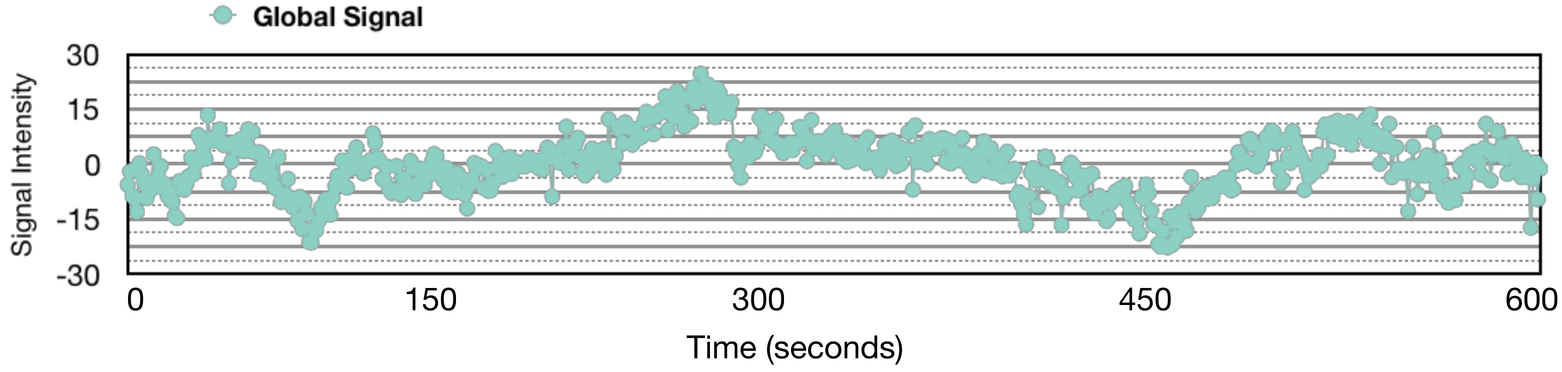


**Supplementary Fig. 3. Physiological monitoring during simultaneous mesoscopic $Ca^{2+}$ and MR-imaging.** Heart rate, breath rate, arterial $O_2$ saturation and body temperature (rectal) are monitored and recorded during imaging experiments. The average and standard deviation of the physiological measurements for each mouse are reported (a.). An example recording from one 10 minute acquisition is plotted (b.). From the same acquisition as shown in (b.), the average MR-signal intensity from the whole brain is plotted in (c.).
