## Supplementary Fig. 5. for "Simultaneous mesoscopic Ca^2+^ imaging and fMRI: Neuroimaging spanning spatiotemporal scales"

### **Supplementary Fig. 5.** Motion within the fMRI time series

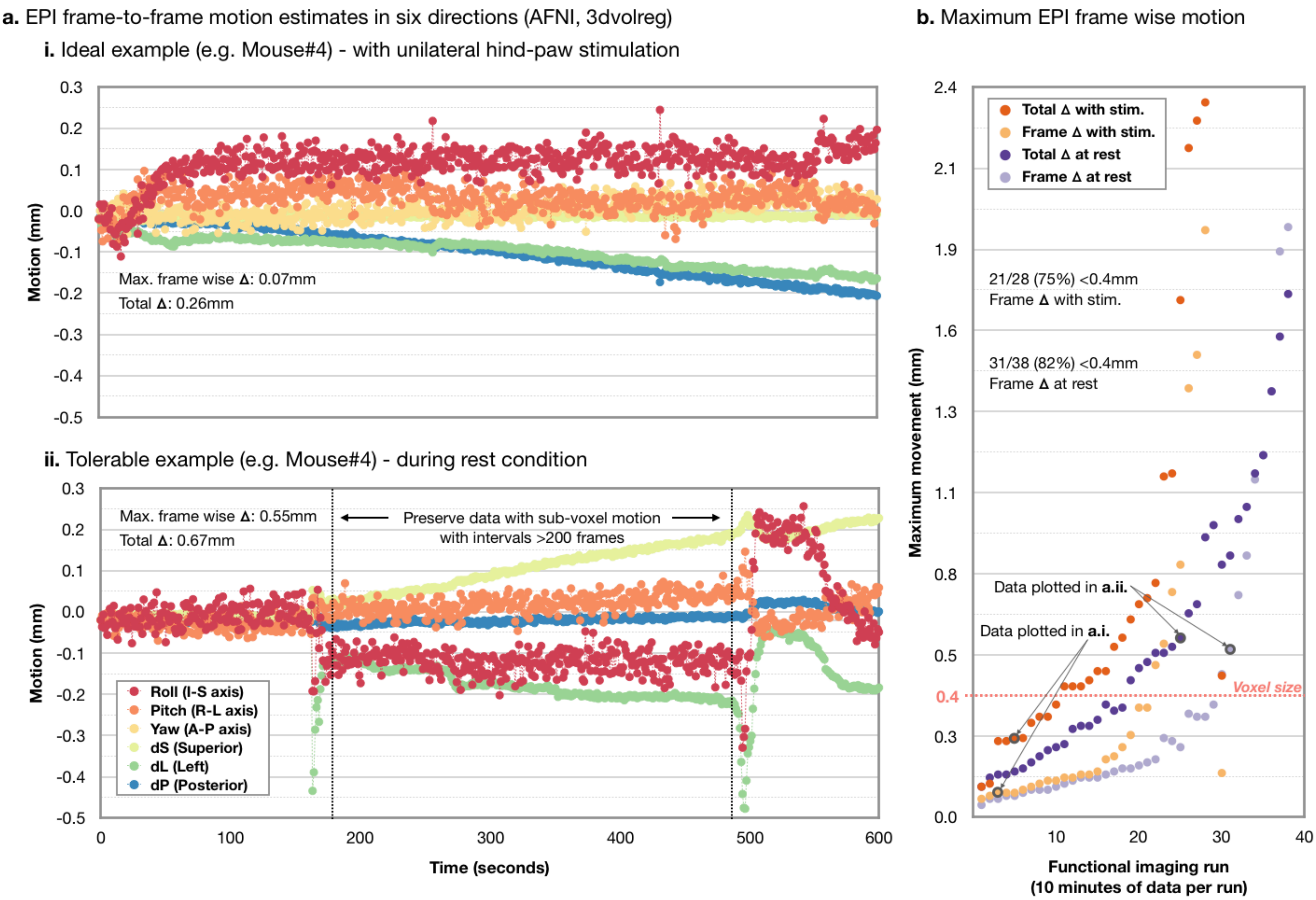


**Supplementary Fig. 5. Motion within the fMRI time series.** For fMRI data, we estimate motion (AFNI, *3dVolReg*) using six parameters (listed in a.ii. Fig. legend). Panel a.i. shows example data from an ideal dual-imaging acquisition. During this experiment, the greatest estimated frame-wise motion was 0.07mm and the total displacement during the 10 minute data collection period was 0.26mm (recall that voxels are $0.4x0.4x0.4mm^3$). Panel a.ii. shows data from a less than ideal example dual-imaging acquisition. During this experiment, there are two instances (denoted by dashed lines) where the frame-wise motion estimate is >0.4mm. However, there is a period between these instances with low motion. Here, 28 acquisitions with unilateral hind-paw stimulation, and 38 rest acquisitions were collected from N=7 animals (each 10 minutes long). In b. the maximum total (dark color) and frame-wise (pale color) displacement estimated during runs with (orange) and without (purple) unilateral hind-limb stimulation are plotted. The examples shown (in a.) are indicated. For the majority of acquisition (75% with stimulation, and 82% without stimulation), the frame-wise motion estimates were <0.4mm.
