## Supplementary Fig. 6. for "Simultaneous mesoscopic Ca^2+^ imaging and fMRI: Neuroimaging spanning spatiotemporal scales"

**Supplementary Fig. 6.** Ray-casting algorithm to create the MR-angiogram projected surface image for multi-modal image registration

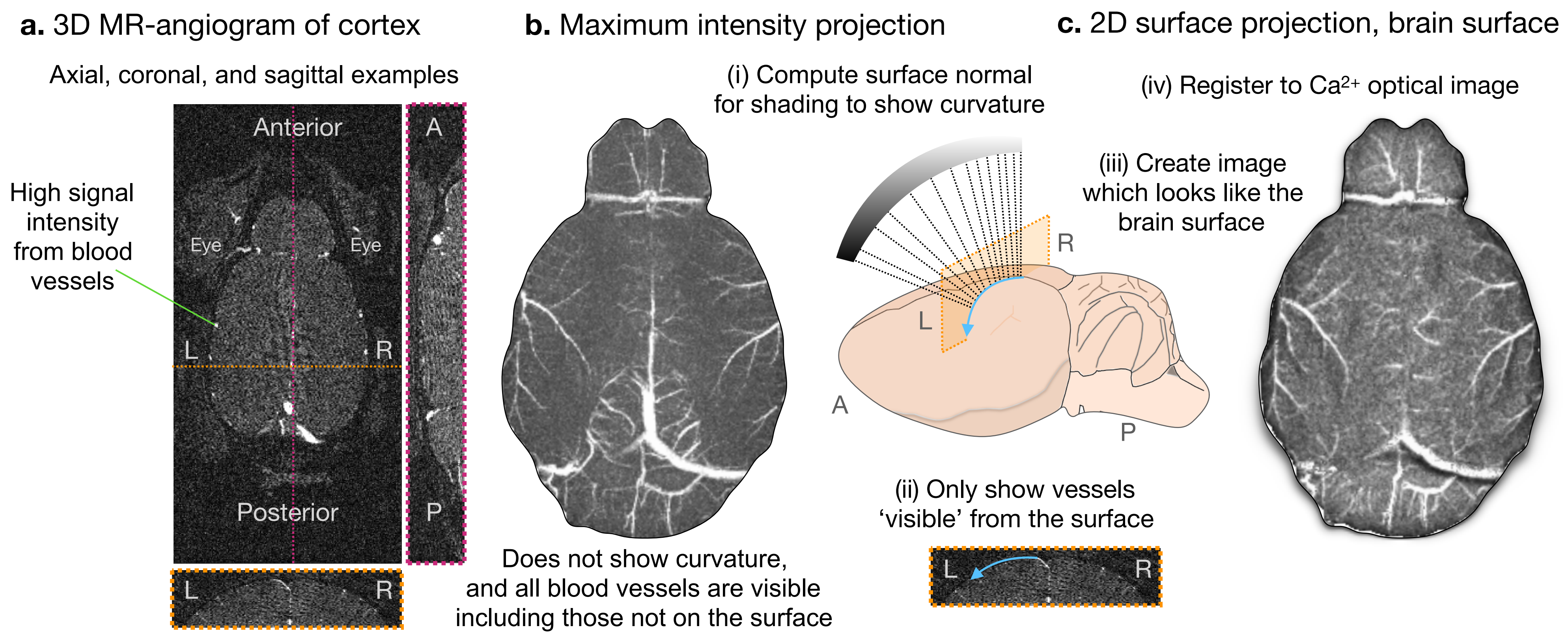


**Supplementary Fig. 6. Ray-casting algorithm to create the MR-angiogram projected surface image for multi-modal image registration.** Three example views of the raw 3D MR-angiogram data are shown (a.). In these images, blood vessels have high MR-signal intensity. Following masking (to remove signal from anatomy outside of the brain) a maximum intensity projection image (b.) can be generated (MATLAB, *max*). However, this image does not show the curvature of the brain surface, and all the blood vessels (whether or not they are 'visible' from 'above') are shown. To recover depth and a sense of surface curvature, and to limit 'visible' vessels to those on the surface of the brain, we project the MR-data along the axis perpendicular to the optical imaging plane. Each pixel is shaded based on brain curvature (Supplementary Material). The result is a 2D projection of the MR image to yield a view akin to the brain surface (b.).
