## Supplementary Fig. 7. for "Simultaneous mesoscopic Ca^2+^ imaging and fMRI: Neuroimaging spanning spatiotemporal scales"

**Supplementary Fig. 7.** Data used to generate the hemodynamic response function (HRF) for the generalized linear model (GLM)

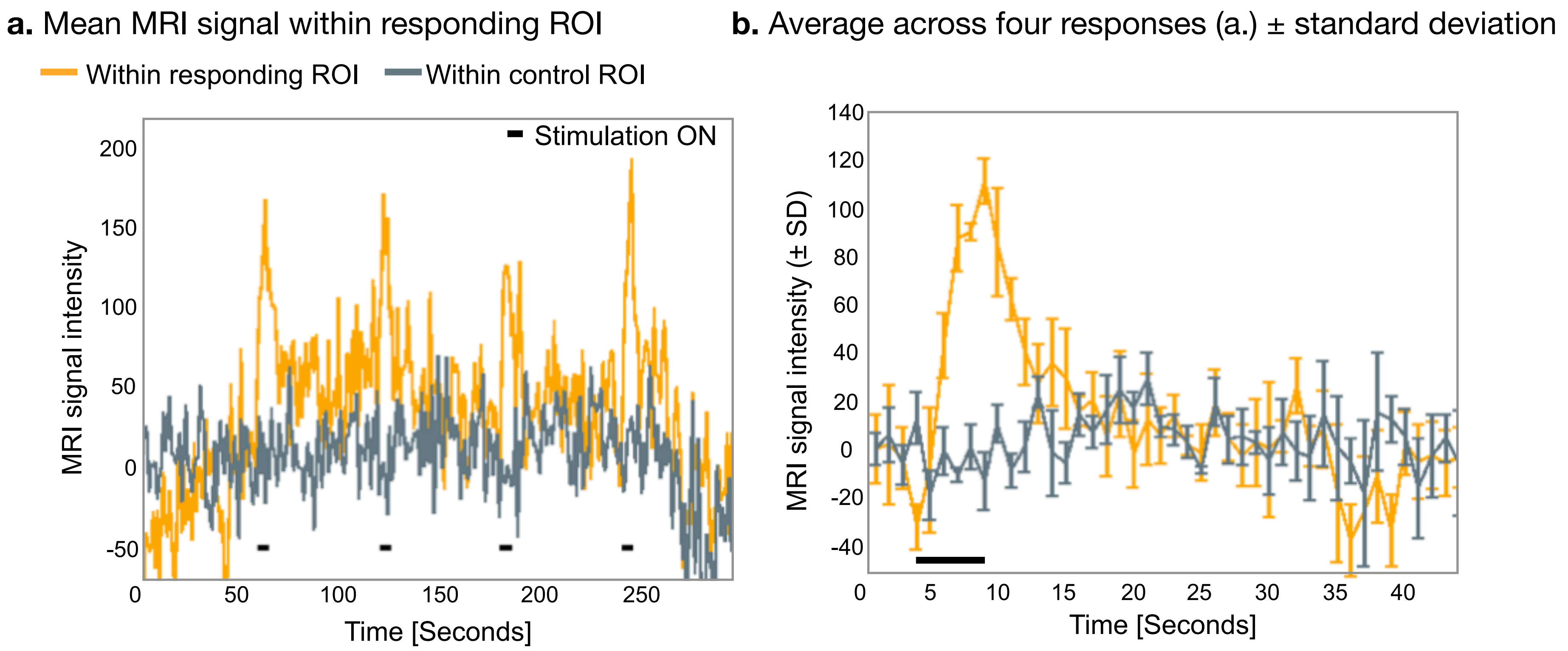


**Supplementary Fig. 7. Data used to generate the hemodynamic response function (HRF) for the generalized linear model (GLM).** A seventh mouse (from a pilot experiment) was used to generate the custom HRF used in the GLM for the analyses of the fMRI data included in Results (N=6). This animal underwent the same imaging protocol (unilateral hind-paw stimulation) as the animals in presented in Results and the data were pre-processed in the same way. To identify the responding ROI, we used a GLM which included the same parameters as the GLM used in Results with one exception: a box-car (in place of an HRF) time-locked to stimulus onset. The average signal within the responding ROI to the first four presentations of the stimulus is plotted (yellow) in (a.). For reference, the average signal within a ROI of the same size located below the cortical strip in the contralateral hemisphere is plotted (grey). The average signal, time-locked to stimulus onset, from the four responses (plotted in a.) is plotted (b.). A smoothed (MATLAB, *smooth*) normalized version of this average signal time-locked to stimulus onset was used as the customized HRF in Results.
