## Supplementary Fig. 8. for "Simultaneous mesoscopic Ca^2+^ imaging and fMRI: Neuroimaging spanning spatiotemporal scales"

### **Supplementary Fig. 8.** Localization of Ca2+ and fMRI responses to stimuli

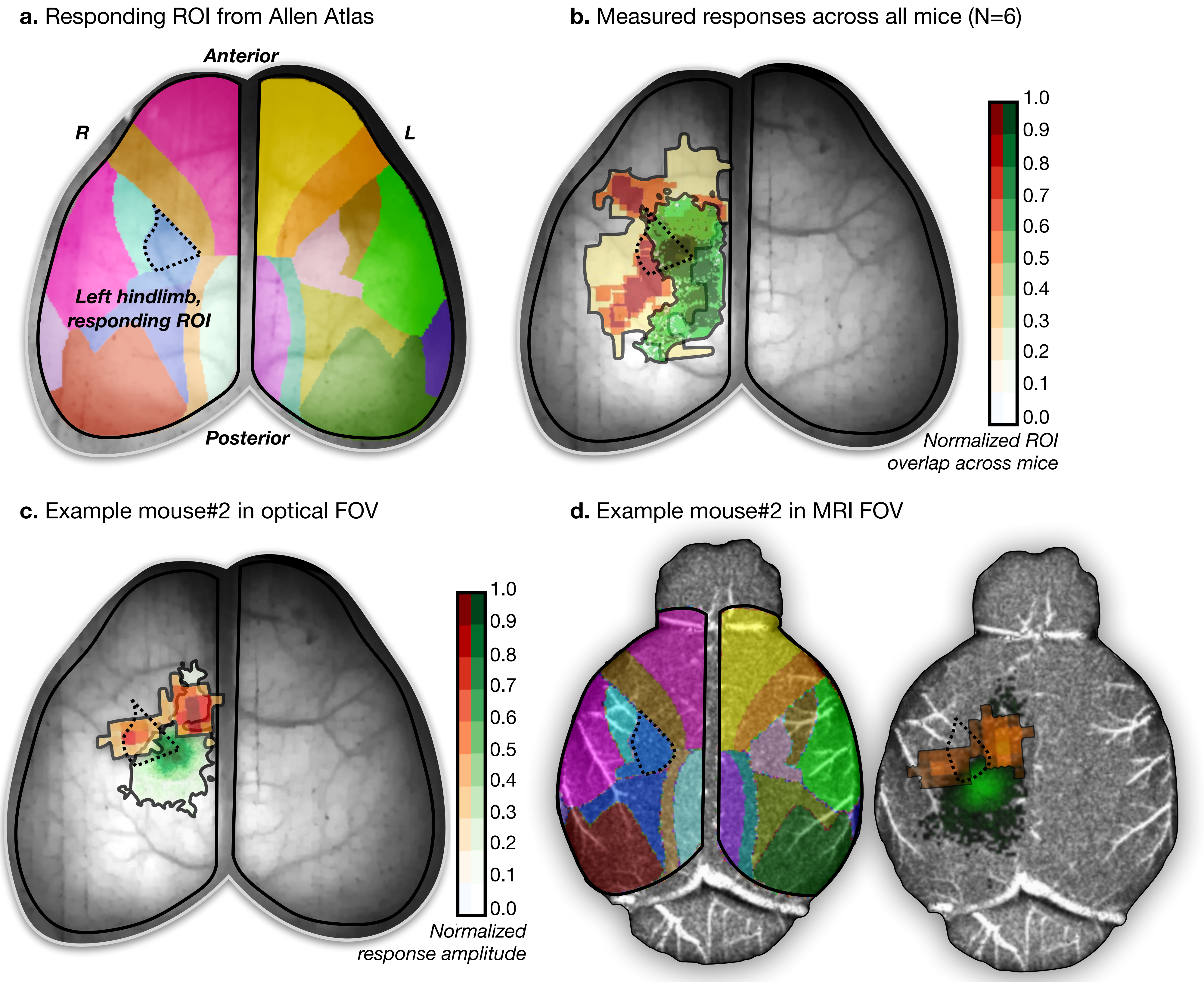


**Supplementary Fig. 8. Localization of $Ca^{2+}$ and fMRI responses to stimuli.** A surface projection of the Allen atlas was registered to all mice. An example of the overlaid atlas is shown in (a.) on the optical data of an example mouse. The ROI expected to respond to the presented hind limb stimuli is indicated with a dotted line. The data from all mice were registered to a common space (the native space of mouse#1). The responding ROIs from all mice from both modalities were normalized to the maximum response amplitude and summed across mice. Overlaid on a representative optical image, the overlapping responding ROIs across mice are shown in (b.). The expected responding ROI from the Allen atlas is shown as a dotted line. An example from one mouse (#2) is shown in (c.). In (d.), the Allan atlas and the responding $Ca^{2+}$ and fMRI ROIs are shown from example mouse #2 (in c.) overlaid on the projected MRI data for this mouse.
