## Supplementary Fig. 9. for "Simultaneous mesoscopic Ca^2+^ imaging and fMRI: Neuroimaging spanning spatiotemporal scales"

### **Supplementary Fig. 9.** $Ca^{2+}$ and BOLD time-to-peak measurement

**a.** Mean $Ca^{2+}$ ΔF/F evoked response

**b.** Mean BOLD evoked response

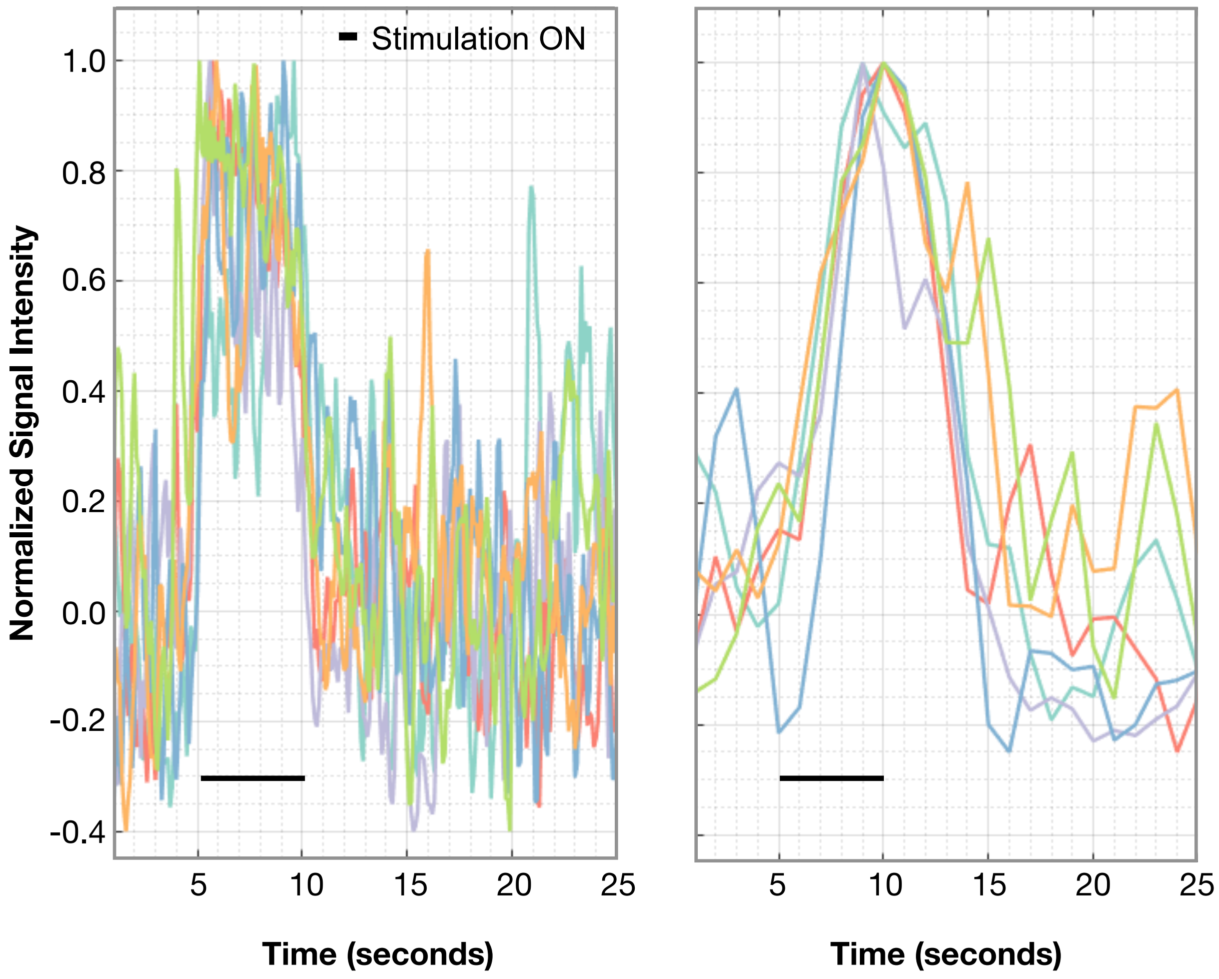


**c.** Time-to-peak summary

| | $Ca^{2+}$ | BOLD |
|---|---|---|
| *Mouse #1* | 0.5 sec. | 4 sec. |
| *Mouse #2* | 0.7 sec. | 5 sec. |
| *Mouse #3* | 0.6 sec. | 4 sec. |
| *Mouse #4* | 0.8 sec. | 5 sec. |
| *Mouse #5* | 0.9 sec. | 5 sec. |
| *Mouse #6* | 0.1 sec. | 5 sec. |
| **Mean ± SD** | **0.6 ± 0.3** | **4.7 ± 0.5** |

**Supplementary Fig. 9. $Ca^{2+}$ and BOLD time-to-peak measurement.** From each mouse (N=6) the time-to-peak response amplitude following unilateral hind-paw stimulation was estimated by averaging across N=9 stimulus responses. Plotted in (a.) is the average normalized (to peak amplitude) $Ca^{2+}$ response for each mouse. Plotted in (b.) is the average normalized (to peak amplitude) fMRI response for each mouse. In (c.), the time of the peak response for each mouse (estimated from (a.) and (b.) is listed with the mean and standard deviation (SD) across mice shown.
