## Supplementary Fig. 10. for "Simultaneous mesoscopic Ca^2+^ imaging and fMRI: Neuroimaging spanning spatiotemporal scales"

**Supplementary Fig. 10.** Parcellation by applying multi-graph k-way clustering

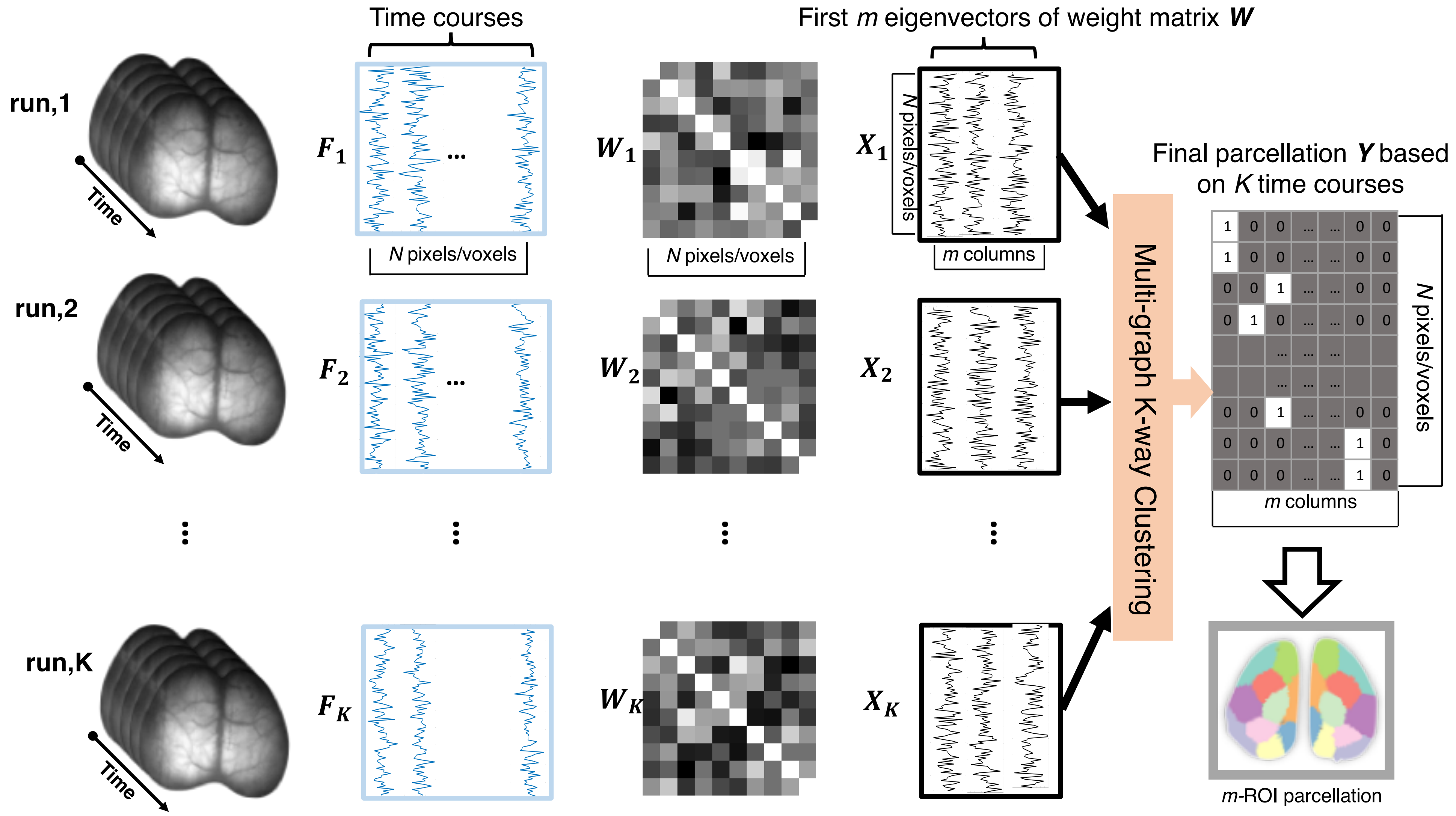


**Supplementary Fig. 10. Parcellation by applying multi-graph k-way clustering.** Individual data sets (from one animal in this example) are shown as runs 1 through 'K' (first column). From each run, we obtain a time-course for each pixel or voxel which is placed within a matrix: 'F' (second column, blue). In the third column, matrices of weights 'W' (NxN) for each matrix 'F' are shown which contain entries that depend on the Pearson correlation between time courses of each paired pixel/voxel (each pair of columns in each matrix 'F'). From 'W', the first 'm' eigenvectors (where 'm' is the *a priori* chosen number of parcellations) are computed ('X', Nxm). These 'X' matrices are passed to the multi-graph k-way clustering algorithm which iteratively solves the optimization problem in the spectral domain (Methods). The solution 'Y' (final column) is converted to a 1D label (vector of zeros and a single one) which is mapped to the original data space (the m-ROI parcellation). Each colored area belongs to a different parcel.
