## Supplementary Fig. 11. for "Simultaneous mesoscopic Ca^2+^ imaging and fMRI: Neuroimaging spanning spatiotemporal scales"

**Supplementary Fig. 11.** Stability of connectivity using $Ca^{2+}$ and fMRI data parcellation

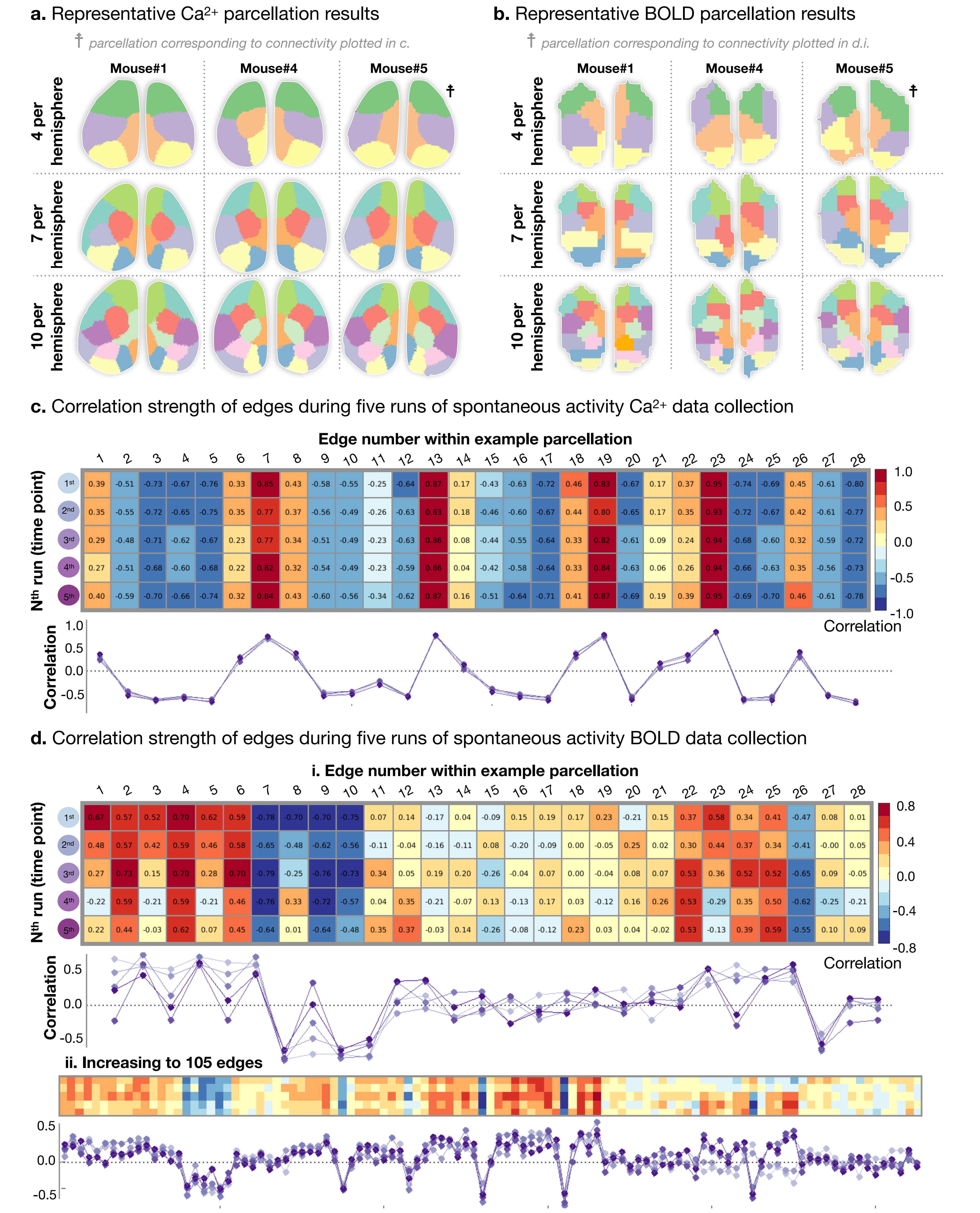


**Supplementary Fig. 11. Stability of connectivity using $Ca^{2+}$ and fMRI data parcellation.** Using multi-graph k-way clustering (Methods), $Ca^{2+}$ (a.) and fMRI (b.) data are parcellated. Three example animals choosing the number of parcels ('m') to be either 4, 7 or 10 per hemisphere. Data from each hemisphere is analyzed separately. Thus, symmetry between hemispheres indicates that the algorithm finds the expected bilateral pattern. Furthermore, similarity between parcellations across modalities indicates the presence of shared spatial features between modalities. This is examined explicitly in Results. Finally, survival of boarders (by the subdivision of large into small parcels) as the number of parcels increases (from 4 to 7 to 10) - in place of the redrawing of new parcels across boarders - indicates reproducibility. In (b.) and (c.) we compute the stability of connectivity patterns throughout the experiment. Ten minutes of spontaneous activity is recorded at five time points which span the imaging protocol. We calculate connectivity between each pair of regions (i.e. connectivity strength along edges) using Pearson correlation. The complete set of edge strengths can be expressed as a vector. For one $Ca^{2+}$ (c.) and two fMRI (d.) example parcellations, we show these vectors assembled into matrices of edges by time points. Below each matrix, a scatterplot of connectivity strength by edge for each time point (purple). The connectivity profile (the pattern of high and low edge strengths) is preserved across time. For 8-10 parcels per hemisphere, the connectivity stability, measured as the Pearson correlation of the connectivity matrix converted into vector format, is very high: $Ca^{2+}$ r=0.995 ± 0.003, and fMRI r=0.84 ± 0.09.
