## Supplementary Fig. 12. for "Simultaneous mesoscopic Ca^2+^ imaging and fMRI: Neuroimaging spanning spatiotemporal scales"

### **Supplementary Fig. 12.** Allen atlas isocortex ROIs compared to data driven parcellation

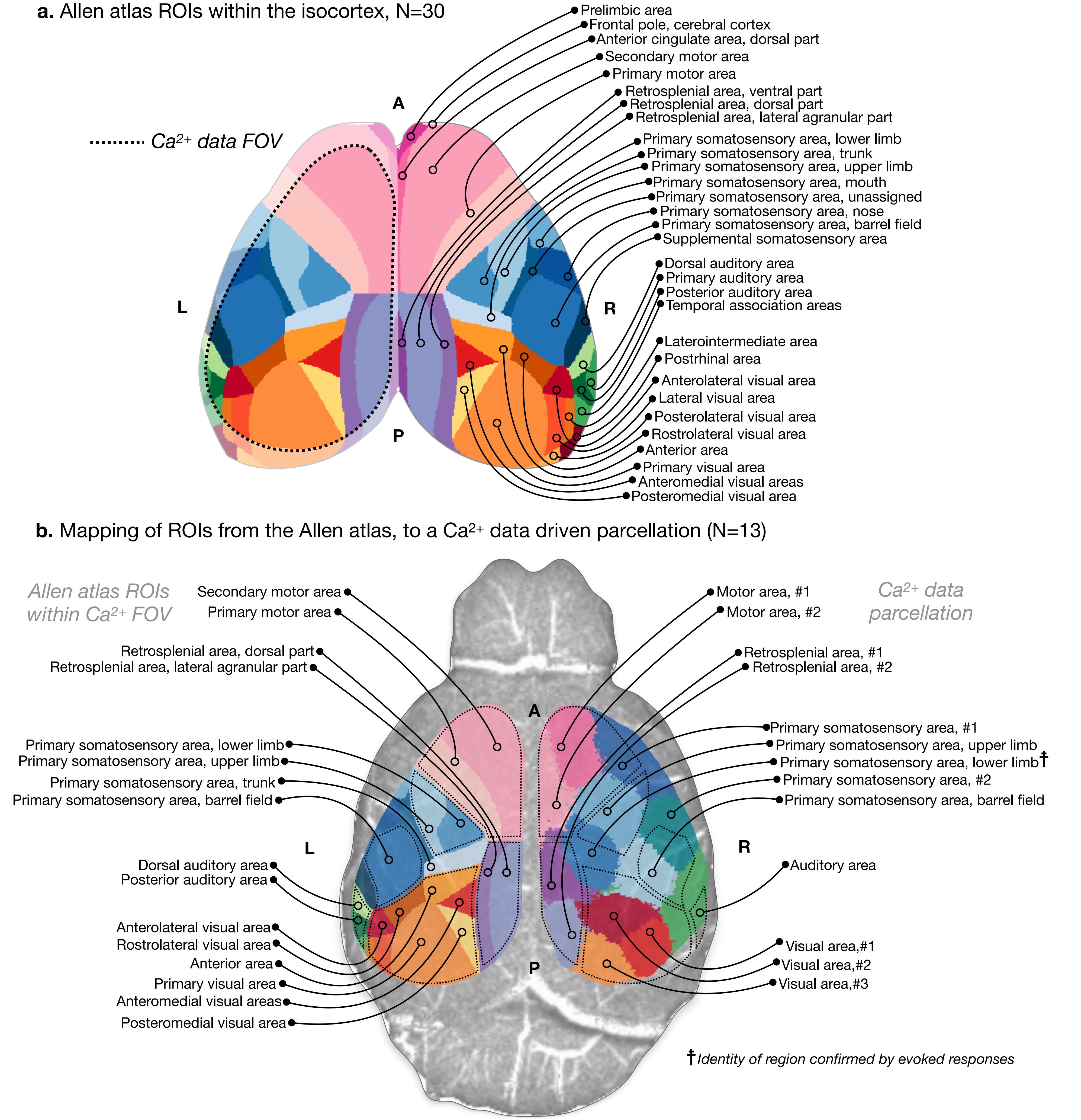


**Supplementary Fig. 12. Allen atlas isocortex ROIs compared to data driven parcellation.** To estimate which ROIs appear within the data driven functional parcellation, we begin with the Allen atlas ROIs that reside within the isocortex (N=30). In (a.) the Allen atlas isocortex ROIs are projected to the brain surface. The dotted line indicates the FOV of the $Ca^{2+}$ data acquisition. In (b.) the Allen atlas isocortex ROIs within the $Ca^{2+}$ FOV are shown overlaid on the left hemisphere grouped by function (e.g. visual areas are shown in orange-red). Functional regions are outlined with dotted lines. A representative functional parcellation (using $Ca^{2+}$ data) containing N=13 parcels is shown on the right hemisphere. The mirror image of the grouped functional areas from the Allan atlas are shown as dotted lines on the right hemisphere. We adopt a naming convention of the functional parcels based on proximity to the Allan atlas functional areas. Note, the naming convention is based on visual inspection with the exception of the lower limb somatosensory area which we identify reliably based on evoked responses.
