## Supplementary Material for "Simultaneous mesoscopic Ca^2+^ imaging and fMRI: Neuroimaging spanning spatiotemporal scales"

*Ray-casting algorithm for MR surface projection image*

Fundamentally, the present algorithm uses the 3D MR-angiogram (**Supplementary Fig. 6.a.**) to create a simulated 2D MR-image (**Supplementary Fig. 6.c.**) that has the same appearance as if the MR-volume were imaged from above by a camera. There are three steps. Step (1) Non-brain tissue from the MR-data is removed using standard mathematical morphology tools (e.g. signal intensity thresholding). Step (2) The MR-volume is rotated and translated using five degrees of freedom: three rotation angles of the axial plane of the MR-volume relative to the optical imaging plane and two in-plane translations. Step (3) The MR-data is projected along the translation perpendicular to the optical imaging plane by shooting ‘rays’ from ‘above’. Each ray is followed until it ‘hits’ the MR brain-volume ‘surface’. For each pixel in the MR-image, we compute the average signal of the first five voxels beneath the surface in the MR-volume and multiply this value by the cosine between the ray and the brain surface normal which is estimated using the local image gradient (**Supplementary Fig. 7.b.**). This generates local shading (the appearance of curvature). Steps (2) & (3) are repeated iteratively. Following each iteration, the similarity of the blood vessels in the optical image and projected MR-image are assessed and the five parameters estimated in step (2) adjusted until the two images converge.

*Custom hemodynamic response function (HRF) from pilot data*

The HRF passed to the generalized line model (GLM) used in the analysis of evoked responses to unilateral hind-paw stimulation was derived from a mouse imaged during a pilot experiment using the same imaging, anesthesia and stimulation protocol as the present work. This mouse was an Ai93 (TIT2L-GCaMP6f, TIGRE - Insulators - TRE2 promoter - LoxP Stop1 LoxP - Green fluorescent protein CalModulin fusion Protein 6 fast), CaMK2a-tTA, Emx1-IRES-cre animal. The strain was purchased from Jackson Labs (JAX stock numbers: 024103, 003010 and 005628) and bred in-house. The Ai93/Camk2a-tTA/Emx1-Cre mice have GCaMP6f expression in excitatory neuronal cell populations just as the Ai93/CaMK2a-tTA/Slc17a7-Cre mice (reported in Results) do.

These fMRI data are processed following the same steps as described in **Methods** (motion correction, blurring, global signal regression, and low-pass filtering), and fit using a GLM (AFNI, *3dDeconvolve*) which includes drift, and motion parameters as described **Methods**. Finally, a threshold is applied to the resulting evoked response map to correct for multiple comparisons (false discovery rate q<0.01) and a cluster size limit (>30 contiguous voxels) applied. The single difference between the analysis of the data reported in Results and the data from this animal is that a box-car time-locked to stimulation onset is used in place of an HRF.

Signal from responding voxels in the contralateral cortex were averaged. We isolate the first four responses to hind-paw stimulation because we find that in a typical experiment the first four responses are the most robust. The averaged, smoothed, and normalized (MATLAB) trace is used as our HRF for the analyses of the N=6 animals presented in Results. The data used to generate the HRF are summarized in **Supplementary Fig. 7**.

*Calculation of the Dice Coefficient*

Given two data sets A and B, the Dice coefficient Dice(A, B) is calculated as Dice(A, B) = 2*|A∩B|/(|A| + |B|), where || denotes the cardinality of the set. When computing the overlap of two parcellations, Parc_A_ and Parc_B_, they are required to be in the same space. Thus, we first find the best matched ROI_B_^i^ in Parc_B_ for a given ROI_A_^i^ in Parc_A_, then we compute the weighted sum of the Dice coefficients as:

$Overlap\left( Parc_{A}, Parc_{B} \right)=(\sum_{i} \left| ROI_{A}^{i} \right|*Dice(ROI_{B}^{i}$, $ROI_{A}^{i}))/\sum_{i} |ROI_{A}^{i}|$
